# Prefrontal cortex temporally multiplexes slow and fast dynamics in value learning and memory

**DOI:** 10.1101/2024.02.02.578632

**Authors:** Seyed Reza Hashemirad, Mojtaba Abbaszadeh, Ali Ghazizadeh

**Affiliations:** Bio-intelligence Research Unit, Sharif Brain Center, Electrical Engineering Department, Sharif University of Technology, Tehran, Iran; School of Cognitive Sciences, Institute for Research in Fundamental Sciences (IPM), Tehran, Iran

**Keywords:** Value, learning, memory, two-rate model, reinforcement learning, single-unit recording, macaque monkeys, prefrontal cortex

## Abstract

Previous studies have revealed segregated circuitries in basal ganglia for fast learning that enables value adaptability and slow forgetting which underlies stable value memories. However, the mechanisms mediating the conflict between value adaptability vs stability remain unknown. Using a reinforcement learning paradigm involving a brief value reversal for objects with previously stable values, we predicted and confirmed a novel behavioral manifestation of the conflict between adaptability vs stability namely the spontaneous recovery of old values in macaque monkeys. Furthermore, we found that individual neurons in ventrolateral prefrontal cortex (vlPFC) temporally multiplexed slow and fast processes in their early and late responses to objects. The local field potential in vlPFC also reflected the two-rate system. These findings implicate vlPFC as a plexus for the interactions between adaptability vs stability in reinforcement learning and suggest spontaneous recovery of past values caused by a two-rate system to mediate relapse to old habits.

## Introduction

Food items, currencies, friends, and sexual partners often maintain stable values over time, but cases arise when values change unexpectedly prompting one to reconsider old reward associations. Consequently, while stable representation of values is shown to be key for skillful interactions with objects such as rapid orienting and ignoring distractors ^1,2^, maintaining flexibility to adapt to new values would be crucial for survival. Nevertheless, the requirements for adaptability vs stability are often at odds. On one hand the system has to learn fast and forget fast if it wants to be adaptable to new values, on the other hand stable value memory requires slow forgetting to maintain old values and slow learning to disregard what can be noise fluctuations in the environment ^3^.

Previous studies have suggested distinct circuits underlying goal-directed (flexible) and habitual (stable) object-reward associations, especially involving the basal ganglia ^4–7^. Specifically, within the macaque striatum, the caudate head (CDh) has been shown to mediate adaptability by fast value learning and fast forgetting, while the caudate tail (CDt) mediates stability by slow value learning but slow value forgetting ^8^. CDh and CDt connect with relatively segregated cortical and subcortical areas and are even enervated by separate groups of dopamine (DA) neurons in substantia nigra pars compacta (SNc) forming separate loops specialized in flexible vs stable reward learning and memory ^9^. However, the interaction between these two systems, in particular when conflicts between past and present reward associations arise, remains unknown. Among various cortical areas, the ventrolateral prefrontal cortex (vlPFC), is a potential candidate to handle the balance between stability vs flexibility in value learning. vlPFC is long recognized for its role in encoding short-term value associated with objects ^10–13^, but more recently its involvement in representing long-term memory of object values is uncovered ^14^. In addition to its multifaceted function, vlPFC is among few areas that take part in both cortical-striatal-thalamocortical loops that control flexibility and stability in reward learning and memory ^15,16^.

Thus, we hypothesized that this cortical area could be a natural hub for the interplay between stable and flexible value encoding. To address this hypothesis, single unit responses and local field potential in vlPFC were recorded while macaque monkeys performed a paradigm in which object values underwent an abrupt reversal in their reward associations. Interestingly, we observed that not only did behavioral output was consistent with a two-rate system with slow and fast dynamics, but that both dynamics were represented within single neurons in vlPFC via time multiplexing.

## Results

A value reversal paradigm was used to study the interaction between stable and flexible object reward associations. Two macaque monkeys (P and H) were first trained to associate abstract fractal objects with preassigned reward contingencies for more than 5 days to create stable object-reward associations ^8,14,17^. Fractals were trained in sets of eight, half of which were associated with high and the other half with low juice amount as reward (good and bad objects, respectively; Fig. 1A). The value training task consisted of force trials where monkeys made a saccade to a single fractal object and received its corresponding reward and interleaving choice trials between a good and a bad fractal to track the degree of value learning (Fig. 1B). The average choice performance was ∼83% at the end of the first day and increased to ∼98.5% after 5 sessions indicating emergence of strong stable learning for consistent with previous reports ^8,18^. Subsequently, the object reward associations underwent an abrupt reversal in one session for a given set such that now the good objects received low reward and bad objects received high reward (Fig. 1D). The monkey’s choice performance following the sudden value reversal, was initially poor but quickly adapted such that they chose the new high value objects after a few trials despite the over-training with the original contingencies (Fig. 1G). The value reversal was followed by a period during which the system was allowed to forget learned values by passage of time and with no additional value learning for those objects (Fig. 1D). This epoch is referred to as the error-clamp period borrowing from a technique previously used to study motor learning and memory ^19^. Note that in this study, we refer to objects as good and bad based on their original overtrained associations with reward before value reversal.

**Figure 1.**
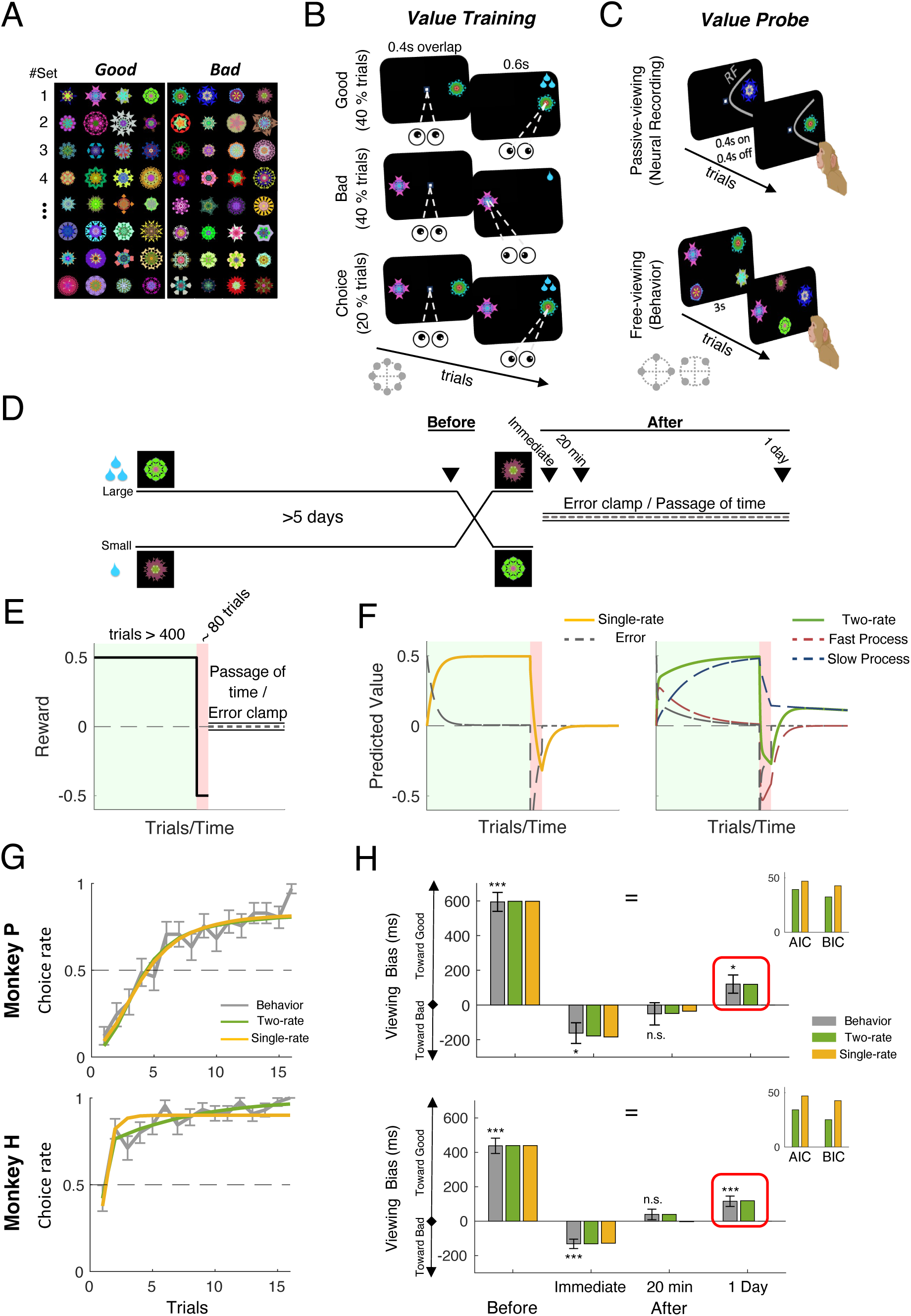
Objects’ value reversal paradigm, model predictions and behavioral results. (A) Abstract fractal objects were associated with reward in sets of eight, four high-valued (good) and four low-valued (bad) (B) Value training was performed with a biased reward association task (forced-choice [FC] task). In forced trials, after central fixation, one object (either good or bad) appeared at one of the eight random locations on screen and monkey had to make a saccade and hold gaze to receive the reward. In choice trials, monkey had to choose between one good and one bad object, appearing at opposite sides of the screen. (C) Value probe was done with passive viewing for neural recording (PV, top) and free viewing for behavioral readout (FV, bottom). In PV monkey had to keep fixating the central white dot while objects (either good or bad) were shown sequentially and randomly in neuron’s receptive field (RF) without reward. In FV trials, four fractals (two good and two bad) were randomly chosen from the set and shown in one of the four corners of an imaginary diamond or square around center. Fractals were displayed for 3s and monkey was free to look at any of the objects or to ignore them. There was no behavioral outcome in this period. (D) Experimental design: A fractal set was initially trained using FC task for > 5 days. First value probe was done by performing PV and FV tasks prior to value reversal. Value reversal was done using FC except that the object reward associations were reversed. This was followed by error clamp period. Three value probes were done during the clamp period immediately after, 20 min after the value reversal and using a set that had value reversal a day before using PV and FV tasks. (E) Reward paradigm of the experiment including the reversal and the error-clamp period. (F) Simulation results of single-rate (left) and two-rate (right) models of the experiment paradigm. Note the reversal of values and reverting back to previous bias (spontaneous recovery) predicted by the two-rate model. (G) Average choice performance of monkey P (top row) and Monkey H (bottom row) during choice trials of value reversal task across sessions (gray line) and predicted choice rate by the best single-rate (orange) and the best two-rate (green) models. (H) Average time difference of viewing good vs bad objects (good minus bad) in FV task across the four value probes and the predicted time difference by the best single-rate (orange) and two-rate (green) models. Inset bar plots show AIC and BIC for the best two-rate and single-rate models. Error bars indicate standard error of mean here and after (SEM). ( = : main effect of memory periods, *p < 0.05, **p < 0.01, ***p < 0.001, here and after).

### Single and two-rate reinforcement learning models

The value learning processes in our reversal task can be modelled using the standard reinforcement learning framework that updates learned values in trial ‘n+1’ as a function of the values in trial ‘n’ modulated by a forgetting rate and a reward prediction error (RPE) arising from the difference between current values and the actual reward that monkey receives multiplied by a learning rate (Eq. 1).

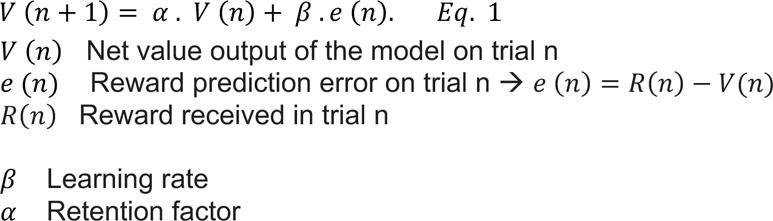

Nevertheless, previous literature on object value learning has revealed two parallel systems in basal ganglia, one that has a slow learning rate and slow forgetting and the other that has a high forgetting and a poor memory but a fast learning rate which is particularly suited for accommodating rapid changes in value such as reversals ^8^. The two-rate system can be modelled by a simple extension of the standard single-rate system by having two processes with different learning and forgetting rates. Given that slow and fast processes in basal ganglia converge on the downstream neurons in superior colliculus ^18^, we proposed the behavioral output to be a simple sum of slow and fast processes (see methods for details).

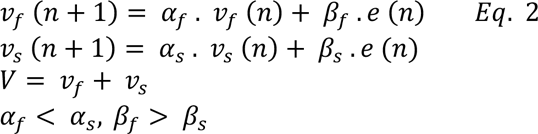

### The spontaneous recovery following reversal and the two-rate model

Figure 1E-F, shows the simulations of the full experimental paradigm and predicted responses in single-rate and two-rate models. In the course of initial training, both models learn the value state of the environment and the error converges to zero. In response to value reversal (sudden rise in error), both single-rate and two-rate predict similar patterns of output reversal which try to track the sudden change in values and sharp rise in RPE. Consistently, both the single-rate (Eq.1) and two-rate models (Eq.2) can reproduce the observed pattern of choice performance going during the reversal task (Fig. 1G, albeit for monkey H the two-rate model had a better fit even after accounting for number of parameters).

Nevertheless, the states of slow and fast processes in the two-rate model show a very different pattern compared to the single-rate model. During the brief reversal, the slow state fails to adapt due to its low learning rate and remains biased toward the initial learning, whereas the fast system rapidly responds and accommodates the value reversal in such a manner that the net value output shifts toward the reversed ones by the end of the reversal session. As a result, two- and single-rate models, can make very different predictions in the error-clamp period. During the error-clamp, the single rate model predicts decay of learned values to zero while the two-rate model predicts spontaneous recovery from the reversed values to the initial values (Fig. 1F). This is because during the error-clamp the fast system forgets rapidly and thus the summed output of both systems reverts back to the initial learned values that is kept by the slow forgetting system.

To test this prediction, the output of the value learning system was probed at four timepoints: before the reversal, immediately after, 20min after and a day after the reversal. Note that special care must be taken such that the value probes minimally affect the learned values themselves and do not cause additional learning. Thus, choice trials similar to those used in value training and reversal are not suited for value probe since they engage the RPE term due to the reward expectation and delivery for each object. Instead, learned values were probed using gaze bias in a free viewing (FV) task (Fig. 1C, see methods). Previous work has shown that free viewing gaze bias is a reliable probe of learned object values ^20^. Importantly, FV task does not engage the RPE term since it is in different context in which there is no reward expectation. As a consequence gaze bias is shown to remain unchanged across consecutive FV trials which would otherwise be reduced if there was any extinction ^21^.

Figure 1H, shows gaze bias in FV in the four time points for both monkeys along with predictions of single- and two-rate models (see methods). As expected prior to the reversal, both monkeys showed a clear bias toward viewing good fractals (monkey H: t (44) = 9.68, P < 0.0001, d = 1.41; monkey P: t (40) = 10.77, P < 0.0001, d = 1.65). Following value reversal, both monkeys’ gaze bias switched toward bad fractals (the new good; monkey H: t (44) = 4.77, P < 0.0001, d = 0.7; monkey P: t (40) = 2.68, P = 0.01, d = 0.4). 20 minutes after reversal, both monkeys gradually forgot the reversal and viewed good and bad objects almost equally (monkey H: t (44) = 1.17, P 0.24, d = 0.18; monkey P: t (40) = 0.7, P 0.44, d = 0.11). Remarkably, a day after the reversal, monkeys’ gaze bias showed a spontaneous recovery toward viewing initially good fractals (monkey H: t (44) = 3.74, P 0.0005, d = 0.56, monkey P: t (40) = 2.26, P 0.02, d = 0.34).

The two-rate model outperforms the single rate model in explaining gaze bias at the four timepoints even after accounting for difference in number of parameters (Fig. S1, best two-rate model AIC < 39.28, best two-rate model BIC < 32.52, best single-rate model AIC < 46.99, best single-rate model BIC < 42.69 Fig 1H inset, see methods). This is because the single-rate model fails to explain the spontaneous recovery rather predicting no bias the day after value reversal.

### Slow and fast processes are multiplexed in early and late firing components in vlPFC

To examine the neural representation of fast and slow processes, acute single unit recordings from vlPFC (121 neurons, area 46v) were performed (Fig. S2). The value probe was done using a passive-viewing (PV) task at the same four time points (Fig. 1D). In PV task monkeys viewed good and bad fractals while fixating centrally without contingent reward (Fig. 1C). The value signal in vlPFC neurons is known to remain unchanged across PV trials as it is a different context from value training task similar to FV making it appropriate for probing value in the absence of RPE ^14^. PV is particularly suited for neural recording since the objects can be shown in the receptive field of a given neuron and are not contaminated with saccadic responses as animals maintain central fixation allowing one to examine purely visually evoked value memories for each object.

Figure 2A shows an example neuron (Monkey H: Neuron # 51) with response time-locked to display onset across the four aforementioned timepoints. This is an example of a ‘’good-preferring’’ neuron that fired stronger in response to the good fractals than bad fractals. Prior to the switch, there was a clear difference in response of the vlPFC neuron to good and bad objects from 100-600ms signaling a well-established value memory in PV. Importantly, following value reversal, the response of the vlPFC was divided into two temporal components. The ‘’early component’’ (∼100-300 ms) of the neural response still showed higher firing to good compared to bad objects consistent with their initial values. But the ‘’late component’’ (∼300-600 ms) showed higher firing to the bad objects consistent with object values after reversal. Interestingly, after 20 min, the late component started to fade away; while the early response was preserved. Since a neuron cannot be kept to test responses a day after reversal, we next showed a set of objects that underwent value reversal a day before to the same neuron, a technique that was used previously to test long-term value memory in neurons ^14,18^. Day-long memory showed preserved preference to the initial values in the early component while there was hardly a response difference in the late component. Figure 2B represents population average response to preferred vs non-preferred object values across value probes. Value preference (good-preferring [Gp] vs bad-preferring [Bp]) of each neuron was determined using cross-validation based on responses before the value reversal (see methods). The population average response clearly showed the presence of early (dark gray) and late (light gray) components. The differential response to preferred vs non-preferred objects, henceforth referred to as the value signal, conspicuously demonstrated the dissociation of these two components across time (Fig. 2B, bottom row). Prior to reversal, neural response was significantly stronger in preferred values (positive value signal) in both early and late epochs denoting the well-learned values (ts (120) > 5.35, ps < 0.0001). Following reversal learning, early response got smaller (F (3,442) = 17.65, p < 0.0001) but remained positive (t (120) > 6.28, p < 0.0001), however, the late response underwent a significant sign change to negative values (t (120) > 5.08, p < 0.0001) as predicted by the two-rate model (Fig. 1F, right). Subsequently over time, the late component decayed toward zero from 20 min after (t (120) = 1.61, p = 0.1) to a day after the reversal (t (82) = 1.03, p = 0.3). On the other hand, the early component was relatively stable and remained significantly positive (after 20 min: t (120) = 5.5, p < 0.0001; next day: t (82) = 7.05, p < 0.0001). Importantly we could predict the average vlPFC responses using the two-rate system and a one-to-one correspondence between early and late components with slow and fast processes, respectively (Fig. 2B, bottom row inset bars). A similar but weaker multiplexing of early and later components in firing was observed during the actual reversal task (Fig. S3a). Note, however that compared to PV results, the responses during the value training are expected to have less power since they are changing over trials due to learning. Furthermore, the firing rate has both visual and saccadic components in this task which can add an interesting yet confounding dimension to the interpretation of neural responses in the active task (Fig S3b).

**Figure 2.**
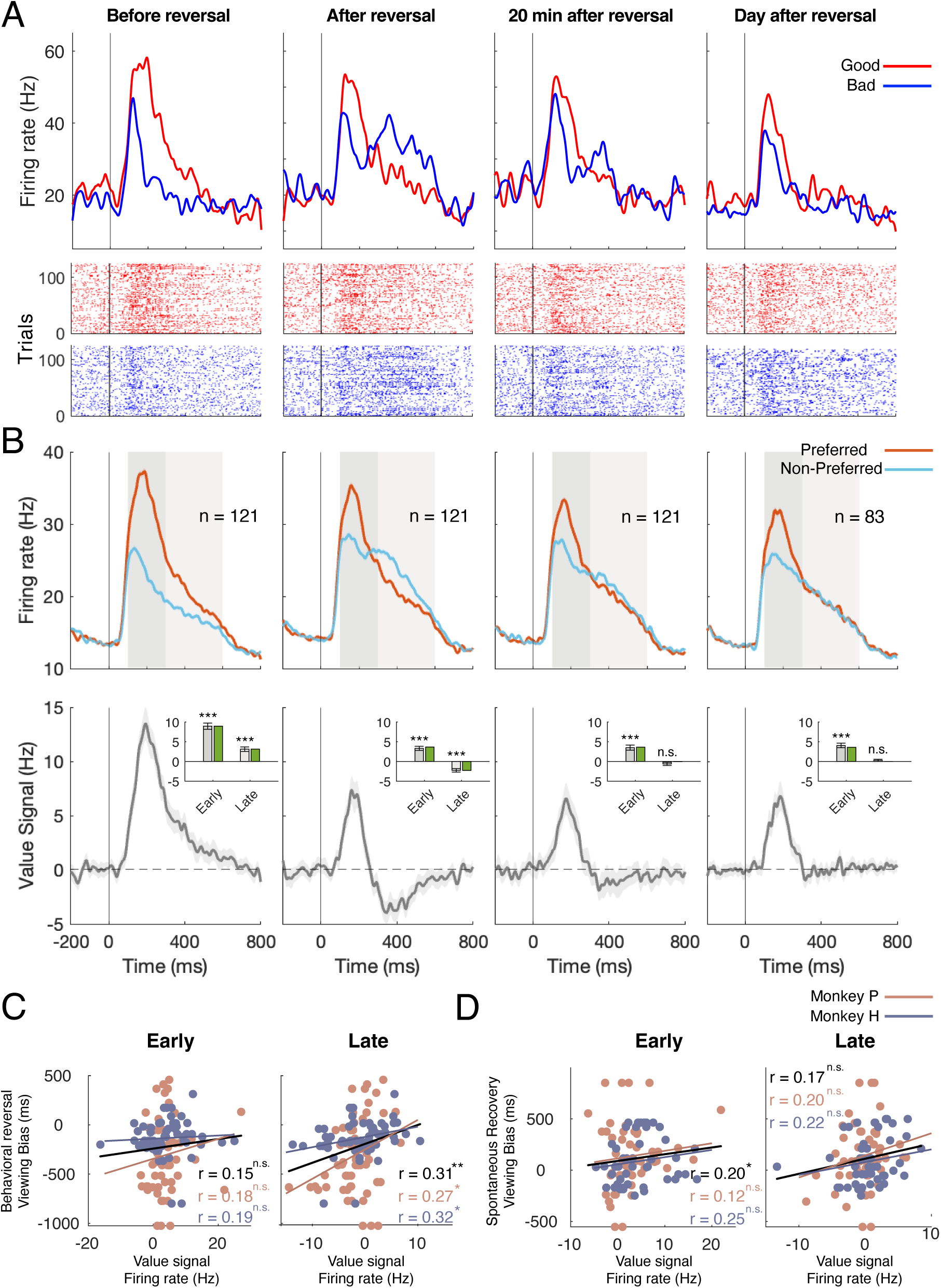
Example neuron and average population response in value probes. (A) The peri-stimulus time histograms (PSTH, first row) and raster plots (second and third rows) time-locked to the display onset of good (red) and bad (blue) objects were shown. The black vertical line indicates the object onset. (B) Average population PSTH for preferred and non-preferred values (top row). Dark and light gray patches represent early (100-300 ms) and late (300-600 ms) epochs respectively. The difference of firing rate to preferred vs non-preferred values (value signal) during experiment (bottom row). The inset bars show the average value signal in early and late epochs along with the predicted values for slow and fast systems by two-rate model, respectively. Shading indicates SEM here and after. (C) Neuron-wise scatterplot of neural value signal for early (left) and late (right) epochs and gaze bias following reversal training. (D) Same format as A but for spontaneous recovery the day after the reversal.

Notably, there was a significant correlation between the behavioral switch in gaze bias following reversal training and the late components of vlPFC neurons (r = 0.31, p = 0.0004), whereas the firing rate of the early epoch and the reversal were not correlated (r = 0.15, p =0.09, Fig. 2C) suggesting that the reversal was mediated by the late component with fast dynamics that adapted to the reversal and not by the early component with slow dynamics which maintained old values. On the other hand, the degree of spontaneous recovery a day later was correlated with the early epoch which survived forgetting (r = 0.2, p=0.03, Fig. 2D) but less so with the late epoch (r = 0.17, p = 0.07). The presence of the unchanging early component in vlPFC firing raises the interesting possibility that the initial saccade in FV should be toward the initially good object even after the reversal. Indeed, this was found to be the case. In both monkeys the first saccade bias remained toward the good objects (Monkey H: t (44) > 2.97, p < 0.0048; monkey P: t (40) > 3.19, p < 0.0028, Fig. S4A) despite the fact that the overall viewing duration clearly switched toward the bad objects after the reversal. This suggests the early component, with slow learning/forgetting, underlies automatic and rapid attentional control toward objects while the fast learning/forgetting process may be involved in the more deliberate later components of attention.

Next, we addressed whether the temporal multiplexing is observed at the level of individual neurons or is mainly a population effect. In the latter case, the early and late components can be segregated across different sub-populations. Furthermore, the stability of the early component may not be a feature of individual neurons but of the population (e.g. as one subpopulation loses the value coding others gain it). These scenarios will have different implications for value coding in vlPFC and the representation of slow and fast processes in this region. Clearly, the example neuron in Figure 2A supports the former possibility. To further address this question the discriminability of good vs bad objects (value signal) across value probes in the early and late component for all neurons was quantified using the area under the receiver operating characteristic curve (termed value AUC here; Fig. 3). Value AUCs significantly above 0.5 indicate higher firing for good compared to bad trials (Gp), and below 0.5 indicates higher firing for bad compared to good trials (Bp). Consistent with previous results of vlPFC, most neurons showed significantly stronger excitation to good objects initially before reversal (AUC_mean = 0.64, t (120) = 15.4, p < 0.0001) in marginal AUC distribution in early epoch and showed little change at various points afterwards (AUC_mean > 0.55, t (120) > 6.78, p < 0.0001). The marginal distribution of before reversal in the late epoch was also significantly biased toward good preference (AUC_mean = 0.56, t (120) = 7.02, p < 0.0001) but immediately after value reversal, the marginal AUC distribution underwent significant change shifting toward bad preference (AUC_mean = 0.45, t (120) = 5.05, p < 0.0001). Notably, 20 min after reversal there was no significant preference (AUC_mean = 0.49, t (120) = 0.23, p = 0.81) nor after a day (AUC_mean = 0.51, t (82) = 1.99, p = 0.049).

**Figure 3.**
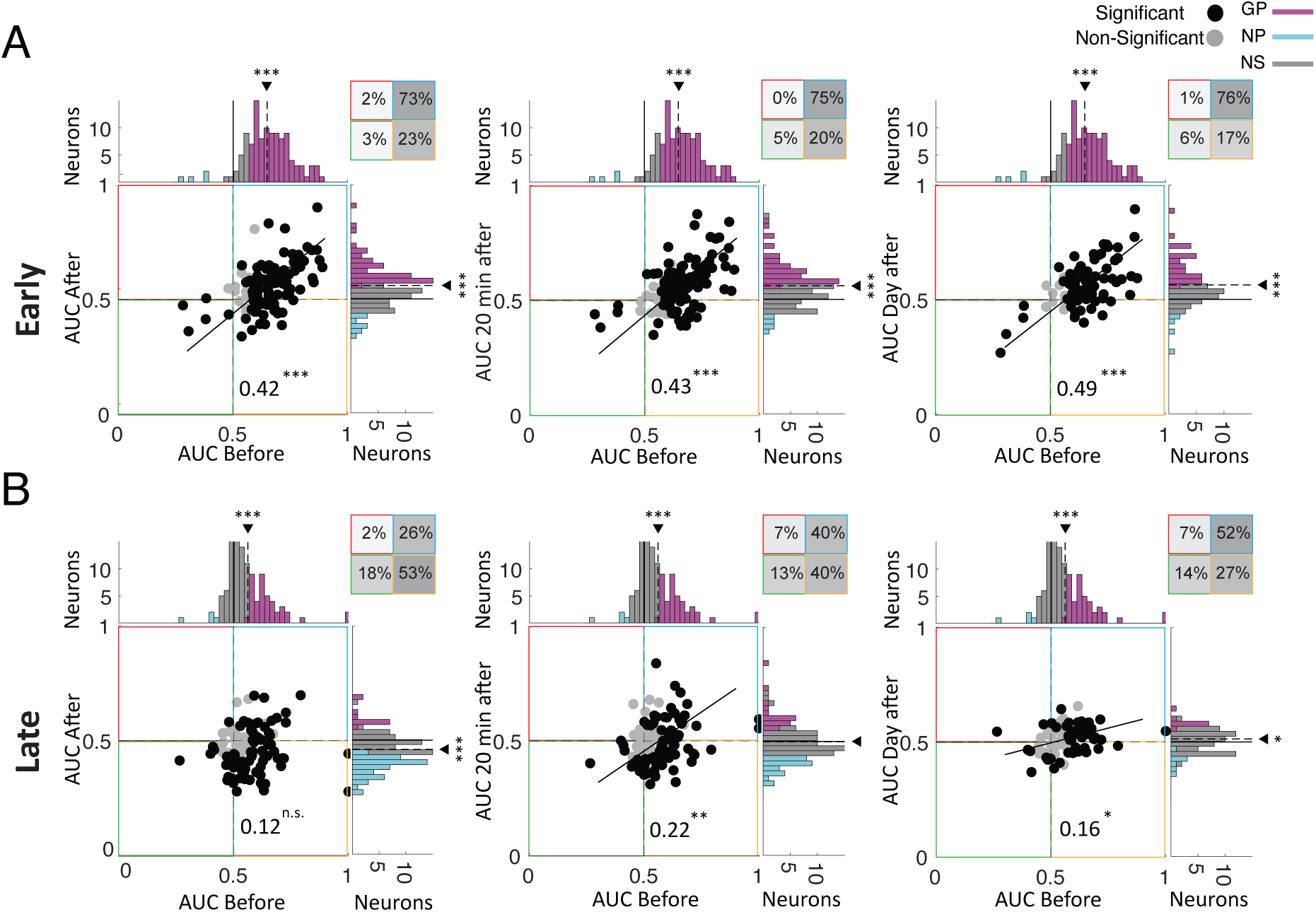
Value responses in individual neurons in early and late epochs across value probes. (A) Scatter plot of pairwise discriminability of good vs bad objects in the early epoch (Value AUC) for immediately after (left), 20 minutes after (middle) and a day after (right) value reversal vs before reversal along with marginal distributions. The magenta, cyan, and gray colors in the value AUC histograms show the good-preferred (GP, significant AUC > 0.5), the bad-preferred (BP, significant AUC < 0.5), and nonsignificant (NS) neurons, respectively. Black dots: significant neurons in either axes; Gray dots: non-significant neurons. The solid black line indicates the linear fit (Deming regression). The inset squares show the percentage of neurons in each quadrant with grayscale color-code. (B) same as A but for the late epoch.

There was a significant positive correlation in value AUC before reversal and all other memory conditions (r > 0.42, ps < 0.0001), indicating a relatively stable value memory in early component across individual neurons. The correlation in value AUC in the late epoch before reversal and after was in general weaker than the early period (p < 0.034, z < 0.069) and was not significant immediately after reversal (immediately after: r = 0.12, p = 0.15, 20 min after: r = 0.22, p = 0.004, day after: r = 0.16, p = 0.017). Importantly, examination of the percentage of neurons with positive or negative value signals (Fig 3A-B inset with <0.5 and >0.5 value AUCs, respectively), showed the stability of value coding in early component (neurons in the first quadrant, 𝜒^#^ = 46, 𝑝 < 0.0001) and value reversal in the late component (neurons in the fourth quadrant, 𝜒^#^ = 16, 𝑝 < 0.0001) following reversal for a significantly large fraction of neurons.

### Population state of the PFC neurons is affected by value reversal beyond the value signal

Previous analysis showed that multiplexing slow and fast processes in the early and late components were evident in a considerable percentage of neurons but there were still some neurons that did not conform to the two-rate model. This suggests that other information maybe encoded by these subpopulations. To get a better view of dominant information present in the population beyond the value signal, we examined the population response dynamics by applying a modified principal component analysis which we refer to as partial-PCA (see methods). Partial-PCA extracts PCA components (in this case the first two dimensions PC 1 & 2) which capture the directions with largest variance orthogonal to the value dimensions that align the most with the value signal present in the neurons (i.e. PCA is done after value dimension is partialled-out). Figure 4 shows the first two PCs plotted against the value axis for early and late responses. As expected, the preferred and non-preferred objects were well-separated along the value axis. Consistent with previous analysis, the polarity of pref/non-pref discrimination remained the same across the four time points in the early component but was reversed in the late component following value reversal. Interestingly, while PC1 showed a trajectory that travelled out and returned a day later to a similar position as before reversal, PC2 showed a sustained displacement from before reversal to a day later (resampling F (1,398) > 19.7, p < 0.0001, MANOVA test). Additionally, there was a significant difference between before-learning and the next day’s neural states (resampling Hotelling’s T^2^ (2,98) > 36.8, p < 0.0001). These results indicate that apart from early and late components of value signal across paradigm, vlPFC population showed irreversible changes in its response due to the experiencing value reversal for the fractals. Using demix-PCA ^22^ also reveals one value probe component which comes back to pre-reversal state and the other that sustains a change even a day later in addition to two value dimensions that roughly correspond to early and late components (Fig. S5). Finally normal PCA also gives qualitatively similar results though less tuned to the relevant dimensions of the tasks as expected (Fig. S6)

**Figure 4.**
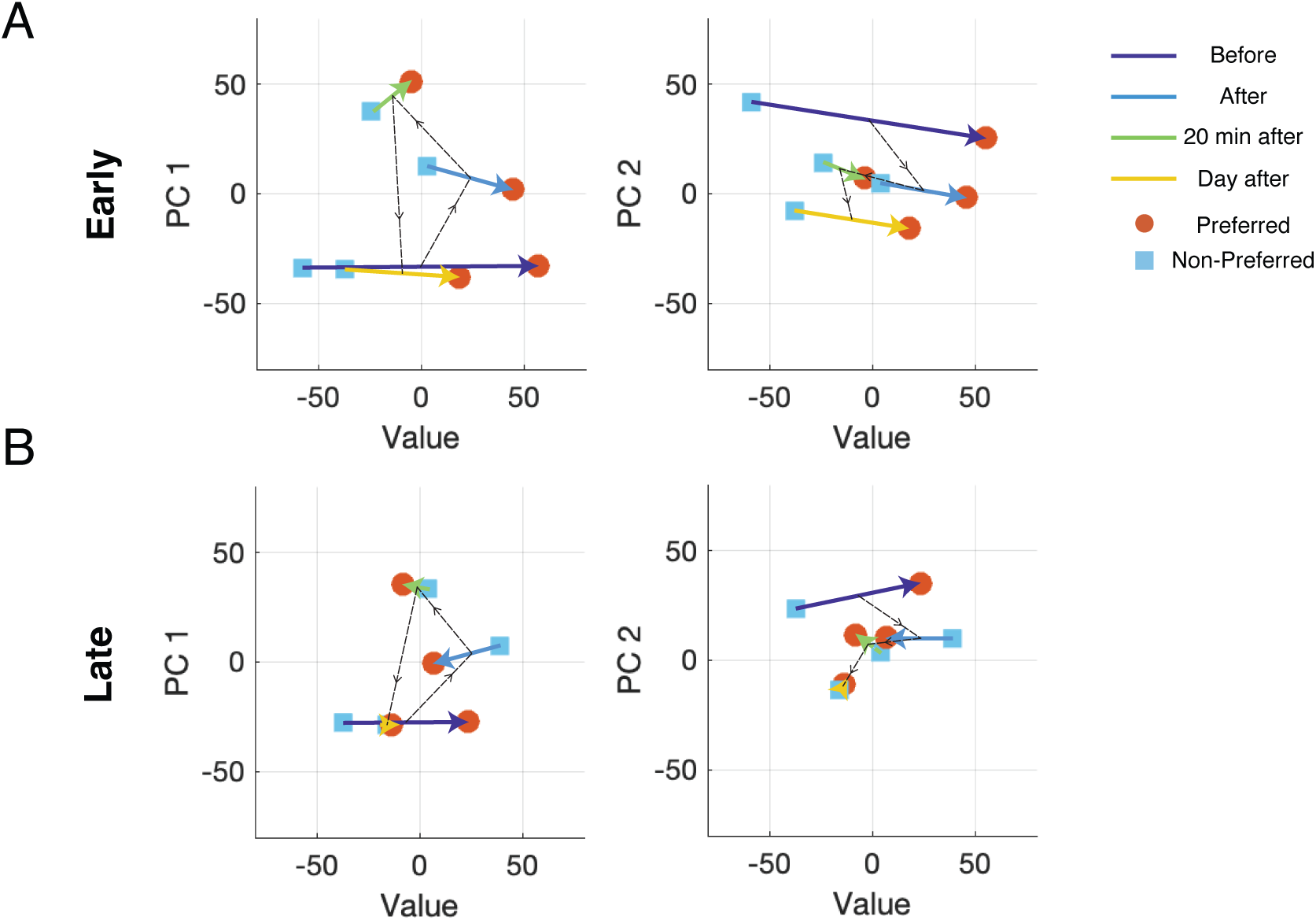
2D projection of population responses using partial-PCA across value probes. (A) PC1 vs value dimension (left) and PC2 vs value dimension for early epoch. (B) same as A but for late epoch.

### The two-rate system is reflected in vlPFC local field potential

Modulations in various frequency bands in local field potential (LFP) are believed to be indicative of local and global computations in neural networks with relevance for dissociating inputs to and outputs from a region ^23–27^. Figure 5A shows the time-frequency power modulations for good and bad objects and their difference (LFP value signal, bottom row) in PV task in the four time points and in different frequency bands. Almost all major band powers including high-Gamma (60-200 Hz), Beta (12-30 Hz), Alpha (8-12 Hz) and Theta (4-7 Hz) showed significant modulation in response to objects and many showed differences for good vs bad object presentations. Among all frequencies the power in high-Gamma resembled the pattern seen in the population average the most with temporal multiplexing of early and late components of its value signal (Fig. 5B). The early component in high-Gamma value signal was significantly positive in all time points including immediately after reversal (t (50-67) > 3.39, ps < 0.001) while the positive value signal in late epoch (t(67) = 7.63, p < 0.0001) became negative following reversal (t(67) = 4.03, p = 0.00014). Similar to the population firing rate the negative value signal in the late component faded away in the 20min after and 1-day later (t(50-67) < 1.85, p > 0.067). On the other hand, there was no evidence of time multiplexing in Alpha and Theta bands as the value signal was similar in both early and late periods after the reversal in Alpha (t (67) = 0.27, p = 0.78) and Theta bands (t (67) = 0.13, p = 0.89). However, both bands showed a strong spontaneous recovery in their value signal (Fig. S7A). In both bands, there was a positive value signal for good vs bad objects (ts (67) > 3.04, ps < 0.003) which was attenuated after reversal (t(67) < 1.56, ps > 0.12) but reemerged in 20min after and in the next day (t(50-67) > 2.28, ps < 0.025). Interestingly, the value signal in 20min after and in the next day was almost as strong as the value signal before reversal (ps > 0.94). The Beta band did not show a significant value signal in any of the time points suggesting that it may not play a role in fast or slow value memories (Fig. S7A).

**Figure 5.**
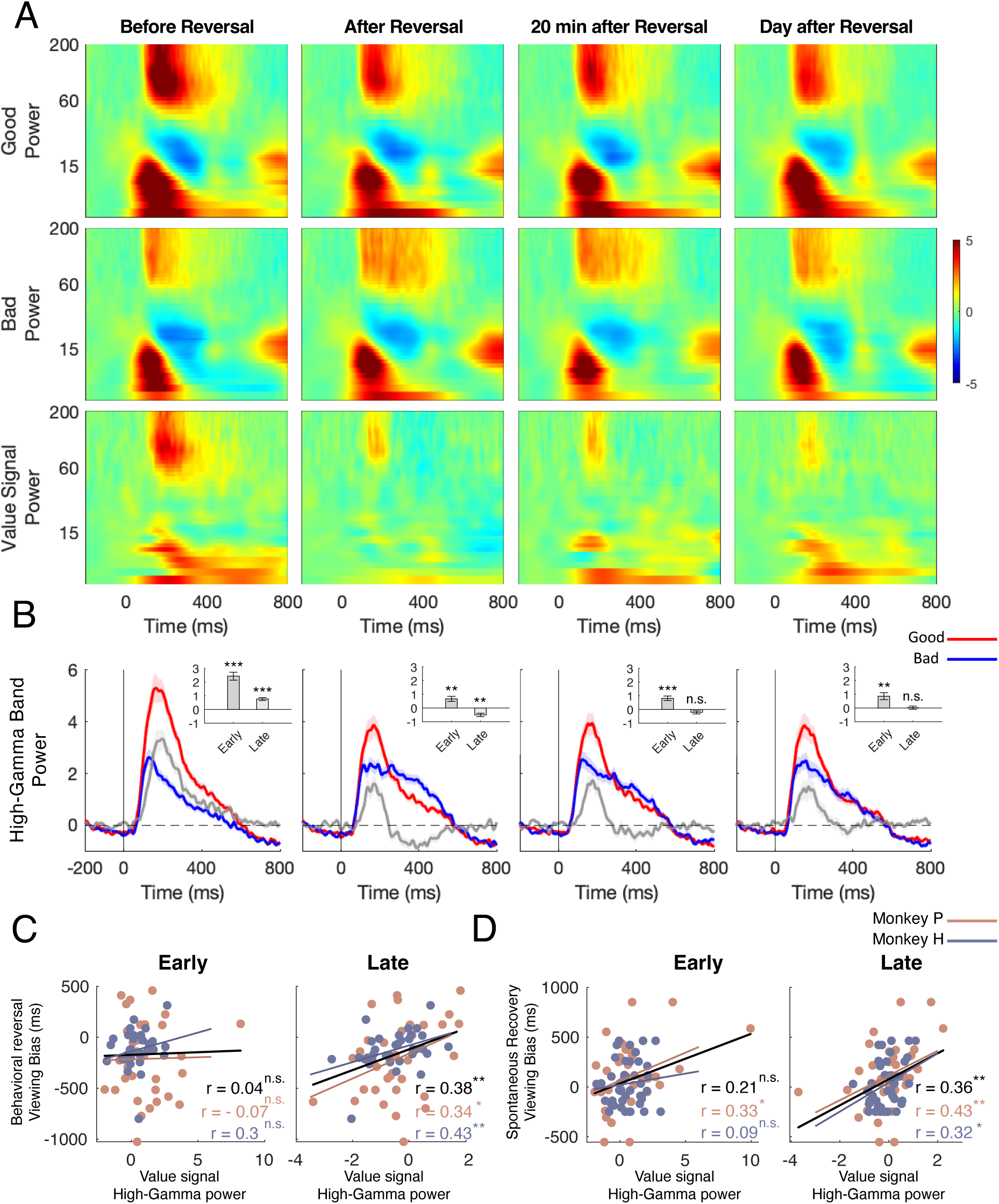
LFP response to good and bad objects across value probes and correlations with behavior. (A) LFP power across time for 0-200 Hz frequencies for good (top row), bad (middle row) and the response difference to good vs bad (LFP value signal, bottom row) across the four timepoints. Colorbar shows LFP power changes relative to (Z-normalized) baseline (−300 to 0 ms). Y-axes are logarithmic. (B) Average high Gamma (60-200 Hz) band power in response to good (red) and bad (blue) fractals and the response difference (value signal, gray). Inset bar plots indicate average value signal in early and late epochs. (C) Session-wise scatterplot of high-Gamma band power value signal for early (left) and late (right) epochs and gaze bias following reversal training. (D) same format as C but for the behavioral spontaneous recovery the day after the reversal.

Similar to what was observed for the firing rates, there was a significant correlation between the late component but not the early component of high-Gamma band power value signal and the behavioral reversal of gaze bias in FV following reversal training (for late r = 0.38, p = 0.001, Fig. 5C). There was no significant correlation between behavioral reversal and neither late nor early response of any of the other frequency bands (only a weak negative correlation between behavioral reversal and the late epoch of Beta band; r = −0.25, p = 0.03; but for others: r < 0.21, ps > 0.07, Fig. S8A). Moreover, a significant correlation was also identified between behavioral spontaneous recovery and the late components of high-Gamma band power value signal and trending correlation with the early component (r _Late_ = 0.36, p = 0.001, r _Early_ = 0.21, p = 0.06, Fig. 5D). Note that unlike reversal which is mainly driven by the fast process in the late component, the spontaneous recovery could show correlation with both components as the fast system forgetting helps the recovery as well. Once again, there was no significant correlation between spontaneous recovery and neither of the other frequency bands (rs < 0.13, ps > 0.24, Fig. S8B). Finally, high-Gamma band value signal showed the strongest and more consistent correlation with the firing rate value signal (Fig. S7B). Interestingly, results showed that there was a decrease in power correlations between the early component of high-Gamma and Alpha/ Theta bands following the value reversal which may suggest a change in the routing of information subsequent to experiencing value reversal (Fig. S7C).

## Discussion

Objects that keep their values stably and for a long time enable rapid and skillful reactions^1^. Nevertheless, changes in old values can happen which requires flexibility to stop automatic reactions and the ability to engage in deliberate interaction based on new values. Here we studied the interplay of slow and fast value learning processes that supported maintaining old values vs adapting to new values in the environment, respectively. Importantly, we showed that a two-rate but not a single-rate system predicted relapse to old values after a period of value reversals a prediction that was then confirmed in behavioral data (Fig. 1). We showed that both slow and fast-learning process were represented in vlPFC population and within individual neurons in the early and late components of their firing (Fig. 2,3). The two-rate processes were also evident in various LFP frequency bands in particular the high-Gamma, Alpha and Theta ranges which showed the effect of reversal and spontaneous recovery to old values (Fig. 5). Finally, there was a significant correlation between value signal in neural firing and in LFP with both the degree of behavioral reversal and spontaneous recovery in free viewing (Fig. 2,5), consistent with the role of vlPFC in the behaviors studied. The slow and fast value learning and memory processes uncovered in this study invokes obvious parallels with the seminal framework in neuroeconomic basis of decision making involving a dual system one being automatic but rapid in its behavioral impact and the other being more flexible but slower to affect behavior ^28^ (Table 1).

**Table 1.**
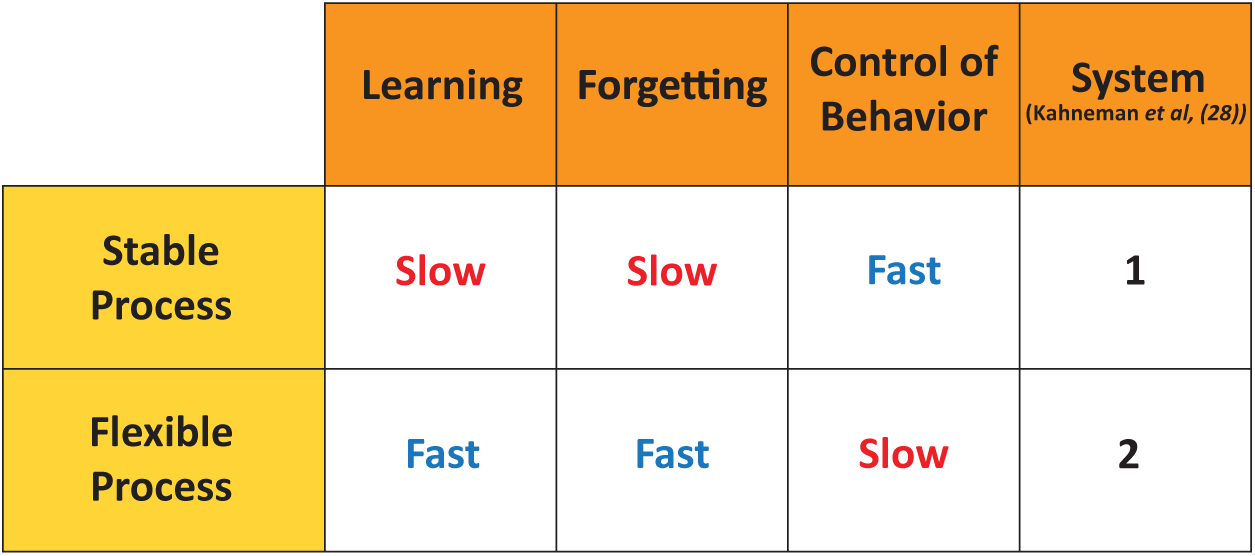
Two-rate processes in the Brain.

|  | Learning | Forgetting | Control of Behavior | System<br>(Kahneman et al, (28)) |
| --- | --- | --- | --- | --- |
| Stable Process | Slow | Slow | Fast | 1 |
| Flexible Process | Fast | Fast | Slow | 2 |

On the behavioral side, our results demonstrated a striking aspect of value adaptation, namely spontaneous recovery of attention to old values despite a period of complete value unlearning (Fig. 1H). We showed that such spontaneous recovery to be diagnostic of a two-rate system rather than a classical single-rate reinforcement model (Fig. 1E,F,H). Such a spontaneous recovery can be a key factor in explaining why old habit die hard and the relapse to old maladaptive behaviors as seen in substance abuse disorders ^29,30^. Such a two-rate system has already been established in motor learning and is shown to successfully explain a slew of phenomena such as spontaneous recovery ^19^, savings (where relearning happens faster than initial learning) ^31,32^, anterograde interference (when learning an initial adaptation slows the learning of an opposite adaptation) ^33,34^. This suggest that a two-rate system might be a fundamental principle underlying adaptation across a wide range of domains from value to sensorimotor learning. Furthermore, such a two-rate system predicts behaviors such as savings and anterograde interference in value learning which is not addressed to-date.

Importantly, neurons in vlPFC reflected signatures of both slow and fast processes multiplexed in the early and late components of their firing rates, respectively. Late response suddenly emerged following value reversal and faded away after ∼20 minutes, signaling flexible value coding, whereas the early component was relatively preserved, indicating stable coding. Interestingly, the early component of vlPFC was comparable to the quick value signal in the CDt appearing ∼100 ms after stimulus onset consistent with a more automatic response to objects. On the other hand, the late response of vlPFC was similar the value signal in CDh with an onset of ∼300ms consistent with a late response to objects ^8^. In addition to the early and late response components, we found sustained changes in the population responses in vlPFC that started from the reversal and remained in the next day. A similar phenomenon has recently been reported in motor learning which involved a wash-out period ^35^.

Our result also showed significant value signal and signatures of the two-rate system in vlPFC LFP. It is generally believed that modulation in different LFP bands contributes to local and long-range network processing with relevance for behavior ^23–27^. In particular, high-Gamma LFPs, originating primarily from interneurons’ network ^26^, had a positive signal correlation with spikes ^25^, suggesting that high-Gamma and spikes are generated within the same network and contribute to the neural output ^25,26^. On the contrary, low-frequency LFP signals can reflect synchronous modulation by the synaptic input ^25,26^ or feedback to the input regions ^36^. Our results regarding high-Gamma power shared the same dynamics as the spiking activity, with the emergence of late response and its abolishment after ∼20 min following reversal, further illustrating that spikes and high-Gamma share the same network of activity. On the other hand, Alpha- and Theta-band showed a single component in value signal that was attenuated following reversal but recovered afterwards. Interestingly, there was a decreased correlation between early components of high-Gamma and Alpha/Theta bands value signal following the reversal which suggests the intriguing possibility that the early component of vlPFC neurons may have become more independent of inputs after stablished value expectations were violated.

In summary, our results showed that value adaptation engages a two-rate system with slow and fast dynamics, that can give rise to the relapse to old habits after a successful washout. Remarkably, our study signifies the role of vlPFC in representing conflicting information by multiplexing stable and flexible values in early and late components of firing. These findings also suggest vlPFC as a novel target for controlling relapse in maladaptive behaviors such as substance use disorders alongside conventional areas in orbitofrontal cortex and ventral striatum ^37–40^. Indeed, the vlPFC connectivity with areas, such as the orbitofrontal cortex ^41,42^, anterior cingulate cortex ^41^ and the amygdala ^43^, which mediate reward-based decisions and error monitoring ^44^ and with pre-supplementary motor area (preSMA) for behavioral switching ^45–47^ as well as directly with SC for saccade control ^15^ situates vlPFC as a central hub to be informed of and manage conflicting reward histories, in particular when the interplay of both short-term and long-term values should be orchestrated for optimal decision making. Future investigations should address the neural mechanisms that allows vlPFC to encode and multiplex early and late value components and whether each component can be independently manipulated.

## Material and Methods

### Subjects and surgery

Two male adult rhesus macaque monkeys (Macaca mulatta) were utilized for this experiment (monkeys P and H, 7 and 12 years old, respectively). All animal care and experimental procedures were carried out in compliance with ethical guidelines established by National Institutes of Health (NIH) and were approved by the local ethics committee at the Institute for Research in Fundamental Sciences (IPM) (protocol number 99/60/1/172). Prior to the experiment, both monkeys underwent surgery under general anesthesia for head holder and recording chamber installment. The head holder was implanted at the midline and a recording chamber was placed and tilted laterally over the right PFC for monkey P and left PFC for monkey H. Following surgery, MRI imaging was performed to confirm the correct position of the chamber for both monkeys. Subsequently, monkeys were trained to learn the experimental tasks after which a second surgery was performed for craniotomy over the PFC region of both monkeys. Neural recording was done through grids with 1-mm spacings placed over the chamber.

### Recording localization

Ventrolateral PFC (vlPFC) localization was performed using T1- and T2- weighted MRI imaging (3T, Prisma Siemens). During imaging, the recording chambers of both monkeys were filled with Gadolinium as contrast agent for enhanced imaging results. Subsequently, AFNI and ImageJ software were exploited to transfer monkeys’ native space into the standard monkey atlas (NMT) to further verify the PFC location and to determine the accessible vlPFC region through recording chamber of each monkey ^48^ (Fig. S2). A total of 62 and 59 neurons were recorded from area 46v ventral to principal sulcus and included in the analysis for monkey P and H, respectively.

### Stimuli

Fractal-geometry objects were used as visual stimuli. Four point-symmetrical polygons overlaid around a common center with smaller polygons in the front comprised a fractal. Size, edges and color of each polygon were chosen randomly. Fractal diameters averaged 4 degrees and were displayed on a CRT monitor. During the course of the experiment each monkey was exposed to a substantial number of fractals (> 500 fractals) in sets of eight for this task and previously published work ^2^. For the current task, monkeys P and H were trained on 44 and 45 sets, respectively which were used during vlPFC recording and/or modelling work presented.

### Task control and Neural recording

All behavioral tasks and recordings were controlled by a customized software program written in C. Neural data acquisition and output control was performed using a Cerebus Blackrock Microsystem device (www.blackrockneurotech.com). Eyelink 1000 plus was used to track eye position during the experiment with a sampling rate of 1 KHz. Diluted apple juice (60 % for monkey H and 50 % for monkey P) was used as reward. During each experimental session, head-fixed monkeys sat in a primate chair and viewed visual stimuli on a 21-inch LG CRT monitor.

Single-unit activity was recorded with tungsten epoxy-coated (FHC, 200 μm thickness) and a laminar microelectrode array (MicroProbes, 16 channels, 125 μm thickness with 250 μm-thickness stainless steel body). For each recording session, the electrode was loaded into a sharpened stainless steel guide tube and then mounted on a Narishige (MO-97A, Japan) oil-driven micromanipulator. The dura was punctured by the guide tube and the electrode was cautiously inserted into the brain with use of the aforementioned micromanipulator. A total of 44 sessions with FHC electrodes for monkey P, 35 sessions of FHC electrodes and 4 sessions of laminar recording for monkey H were recorded.

The electrode’s electrical signal was amplified and filtered (1 Hz to 10 kHz) and subsequently digitized at 30 kHz. Blackrock online sorting with hoops method was used to isolate the unit spike shapes. The online peristimulus time histogram (PSTH) and raster plots of the selected unit were created using a custom code written in MATLAB 2018b (Mathworks inc). All well-isolated and visually responsive neurons were recorded. For the actual analysis of each session, offline sorting with Plexon offline sorter was used afterwards (Plexon, Dallas, TX, USA). In this analysis, first, a negative threshold of 3-3.5 standard deviations away from median were applied to detect spike data. Then, using principal component analysis (PCA) and Plexon built-in template sorting algorithm, the spike data was isolated and clustered into different units. All units for which less than or equal to 0.5 % of the population fired in the refractory period of 1.2 ms were considered as single units. To select visual responsive neurons, custom-written MATLAB functions were used. In short, first, the average PSTH time-locked to display onset across all trials for each neuron was computed. Then, the data was z-scored using the baseline from −200 to 0 ms relative to the object onset. Using MATLAB *findpeaks*, the first response peak after object onset was detected. The minimum threshold for peak height was 1.64, corresponding to the 95% confidence interval. The first valley before the first detected peak was taken as the visual onset of the neuron. Onset of the value signal was also detected using this algorithm on the average value PSTH of individual neurons. All visual responsive neurons that maintained the visual response throughout the paradigm were included in this study.

### Neural data analysis

All neural responses were time-locked to object (visual stimuli) onset. The main analysis epoch was from 100-600 ms after stimulus onset in passive viewing task which was further divided into early (100-300 ms) and late (300-600 ms) epochs. The area under the receiver operating characteristic curve (AUC) was used as a measure of discriminability of objects’ values based on the firing rate across trials in response to good and bad objects in the corresponding epochs. Wilcoxon rank-sum test was used for AUC’s statistical significance for each neuron. The preferred object value for each neuron was determined using a cross-validation technique ^14^. A deconvolution technique was also used to dissociate the visual and saccadic responses in the PSTH of reversal training task (Fig. S3B) ^49^.

### Population analysis

To explore population dynamic changes following reversal, principal component (PCA) analysis, an unsupervised dimensionality reduction method was used. The high dimensional data was the matrix 𝐷 with rows equal to the number of neurons and columns equal to two values (preferred and nonpreferred) by four memory timepoints for each neuron resulting in a total of eight columns. The average firing rate in 100-600 ms across all trials was used to fill the eight columns for each neuron. Subsequently, MATLAB *pca* function was utilized with singular value decomposition (SVD) algorithm to extract components with maximum explained variance. Eventually, the transformation vector was used to project the average neural response in early and late components to PCA states, separately, yielding early and late neural states. The three leading components were plotted (Fig. S6).

The dimensions found by conventional PCA do not necessarily encode the value dimension but find directions with most explained variance. To find the dimension that was most informative about value coding in the region a partial-PCA (pPCA) method was developed and applied to the data. In this algorithm, we first identified the two projection axis that maximized the value signal in the neural data (since the initial value vector did not capture the late value signal, we repeated this process two times).

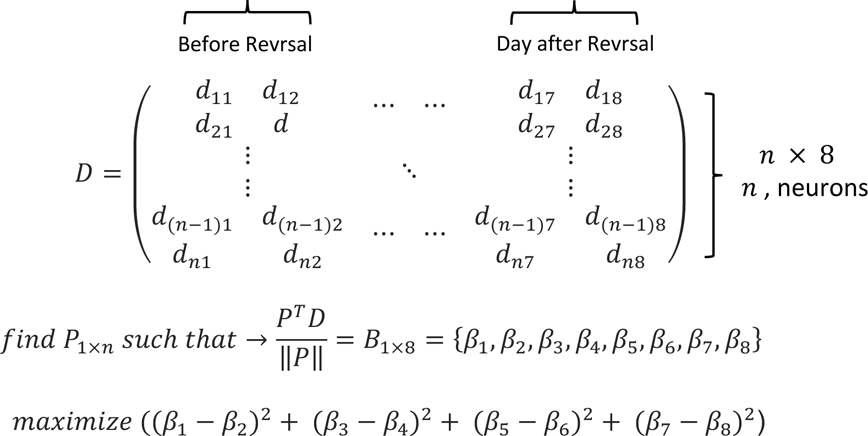

Subsequently, conventional PCA was applied on the residuals to find the remaining directions that explained the most variability in the data. The residuals 𝐷 ′ were calculated according to the following formula:

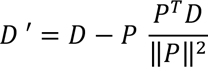

To better visualize the data, since the two value components were orthogonal to each other, we took the sum of the projected values and plotted them against the other two residual PC leading components (Fig. 4). The pPCA results were similar to the results using previously introduced demixed-PCA method by Kobak, et al. ^22^ which addressed a similar limitation in normal PCA (Fig. S5).

### Local field potential (LFP) Analysis

For the LFP analysis, data was down-sampled to 1 kHz and band-pass filtered between 0.1 to 250 Hz with EEGLAB (-v2022.0) FIR filter. Line noise and its sub-harmonics were removed using notch filters. For each session, trials with an LFP amplitude range of 3 standard deviations from the median were identified as noisy trials and removed in pre-processing. The time-frequency analysis was performed using a continuous wavelet transform (CWT) with 7 cycles. For this purpose, the Fourier transform of the analytic Morlet wavelet and the signal was calculated separately. The resulting Fourier transform of the wavelet and the LFP signal were then multiplied following an inverse Fourier transform to obtain the CWT time-frequency results ^50^. Zero-padding was performed in order to avoid edge artifacts. Subsequently, baseline normalization was performed on the resulting time-frequency power, using z-scoring method with the baseline of −300 to 0 ms relative to object onset. In this study, five frequency bands were defined as Theta (4-7 Hz), Alpha (8-12 Hz), Beta (12-30 Hz), Low-Gamma (30-60 Hz), and High-Gamma (60-200 Hz).

### Value Training: Force-Choice (FC) Task

A biased reward association task was used to train object values in monkeys. Each session of training was performed with one set of eight fractals (4 good /4 bad fractals). In each trial, initially, a white dot appeared on the center of the screen (2 degrees) for monkeys to fixate. After maintaining the fixation for 200 ms, either a high or low valued fractal object was displayed in one of eight peripheral locations at 9.5 degrees eccentricity. Following a 400 ms overlap, the central dot disappeared that worked as a cue for the monkey to make a saccade to the fractal. Monkeys were required to hold the gaze for 500 ± 100ms to receive small (low value) or large (high value) reward. A variable inter-trial interval of 1-1.5 s following reward delivery initiated with blank screen. A correct tone was played after a correct trial; however, premature saccade or breaking fixation resulted in playing an error tone. The high to low juice reward ratio was 3 to 1 in both monkeys. Each session consisted of 80 trials, including 64 force trials with the aforementioned design and 16 choice trials pseudo-randomly interleaved between force trials in such order that every 5 trials had one choice trial randomly presented in one of the 5 trials. Choice trials had the same structure as force trials except that one high valued and one low valued object appeared simultaneously on opposite sides of the screen and monkey had to make a choice. Monkeys’ choice rates were used as a measure of objects’ value learning.

### Receptive field (RF) Mapping

In this task, monkeys had to keep fixating on the central white dot (2 degrees) while fractal objects were shown in one of the 32 locations spanning eight radial directions 45 degrees apart and eccentricities from 0 to 11.2 degrees in four steps pseudo-randomly in a manner that all eight fractals of a given set were presented once in each of the locations. Each session consisted of 64 trials. In each trial, four fractals were sequentially presented with 400 ms on and 400 ms off period. Animals were rewarded for fixating after every four object presentations with medium reward following an inter-trial interval of 1-1.5 s with black screen. Subsequently, the online peristimulus time histogram (PSTH) and raster plots of the selected unit time-locked to fractal presentation onset was created using a custom code written in MATLAB 2018b (MathWorks inc) and average firing rate across trials in 100-400 ms window was computed for each location and the maximum response location was used as the receptive field of that neuron (RF-In area) for subsequent passive-viewing tasks in the recording session.

### Neural value memory: Passive viewing task

Passive-viewing task design followed the same structure as the RF mapping task except that the objects were shown close to the location of maximal visual response (RF-In) for each neuron. In case of laminar recording in which more than one channel was involved, the passive-viewing task was performed in all the 32 mentioned locations. Subsequent analysis for receptive field mapping was carried out offline. Analogous to the online mapping the average response across trials in the mentioned window for each location was computed and the locations for which the average response exceeded 70 % of the maximum response was considered RF-In locations and were included in further analyses.

### Behavioral value memory: Free viewing task

Each free-viewing session consisted of 20 trials with one set of fractals. In any given trial, four fractals (2 good and 2 bad) were randomly chosen from the set and shown in one of the four corners of an imaginary diamond or square around center (9.5 degrees away from display center). Fractals were displayed for 3s during which time the monkey was free to look at any of the objects or to ignore them. There was no behavioral outcome in this period. Then the fractals disappeared and after a delay of 0.5 to 0.7 s, a white dot appeared in one of the nine random locations on the screen (center and eight radial locations). Monkey had to make a saccade to the dot to get a medium reward; however, this reward was not contingent with the free viewing period. After reward delivery a black screen with ITI of 1-1.5 s would ensue.

### Eye data analysis

Subsequently in offline analysis, gaze locations were analyzed using custom-written MATLAB functions to extract saccades (displacements > 2.5 degrees/sec) and stationary periods. Initially, trials in which monkeys viewed the screen for less than 1s were removed as a measure of lack of concentration. Then time of viewing the fractals as a behavioral measure of gaze bias was computed. Additionally, we calculated the probability of the first saccade to the initial good objects. Behavioral data of this task was from 44 sets of monkey P and 45 sets of Monkey H.

### Experimental design

For a given set of eight fractals, monkeys were initially trained to associate 4 fractals with high reward (good objects) and 4 fractals with low reward (bad objects) for more than five days to establish a well learned memory of the objects’ values. Monkeys’ choice rate was used as a measure of value learning. In addition, monkeys’ choice performance had to be >90% for two consecutive training days for a set to pass on to the recording session. In the recording session, after isolating a given unit, RF mapping task was done to find the neuron’s receptive field. After that passive-viewing and free-viewing tasks were performed to examine the established neural and behavioral value memory, respectively before value reversal (1^st^ time point). Subsequently, the monkey did a value reversal task. Value reversal task had the same structure and number of trials as a regular FC block except that the reward associations of good and bad objects were reversed (making saccade to good objects resulted in low reward and vice versa). Each set was used only once in the reversal task. Monkeys learned to adapt quickly to this abrupt switch of values and reversed their choices toward the initially bad objects to get the maximum reward. In some sessions monkey P choice rate after reversal remained below 30% which were excluded from further analysis (3 out of 44 sessions in monkey P). All sessions for monkey H were included in the analysis as the correct choice rate was above 30%. Reversal was followed by passive-viewing and free-viewing tasks immediately after (2^nd^ time point), 15-20 minutes after (3^rd^ time point) and a day after the reversal (4^th^ time point) to probe changes in neural value memory and behavioral gaze bias, respectively. Monkeys engaged in a search task with a different set of fractals in the intervening time between the 2^nd^ and 3^rd^ time points to keep them engaged with different objects ^2^. Since it is not normally possible to keep the same neuron across days the response of the neuron on the 4^th^ time-point was probed with another set of fractals that underwent reversal a day before. A similar method was used by us previously to look at the longevity of value memory across many days in a single neuron ^14^. This method is especially suited for vlPFC as its neurons are shown not have a large object selectivity to interfere with their value coding ^14^.

### Modeling

In this study, two models were used to outline the progression of value memory throughout the course of the paradigm: a single-rate model and a two-rate model. In both models, value learning in a particular trial is a function of current value state and reward prediction error (RPE) experienced in that trial. The learning rule for these models are as follows:

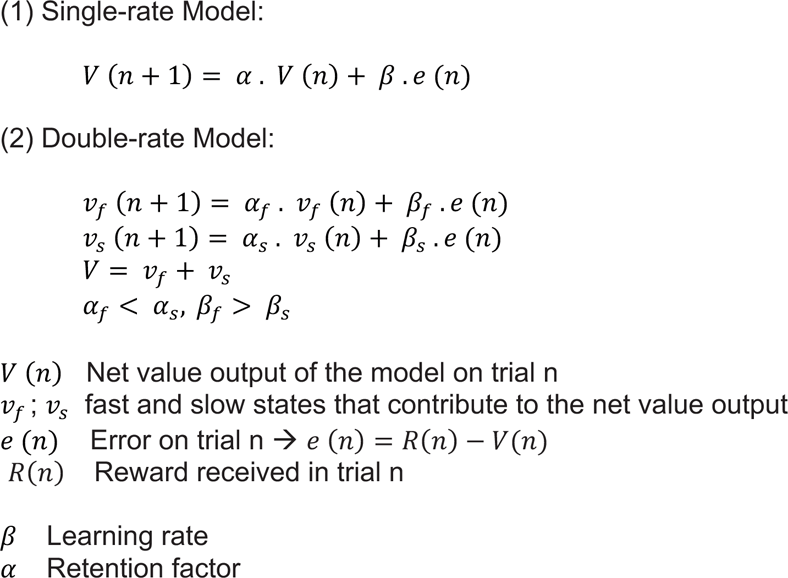

Where ‘’n’’ is the trial number. The error arises because of the difference in predicted value by the model and the reward monkey actually receives. In the single rate model one retention and learning rate was used, whereas the two-rate model consisted of two interactive systems each with their own retention and learning rates (equations 1 and 2). In the two-rate system we assumed that one system has a higher learning and lower retention rate (i.e. higher forgetting rate) which we refer to as the fast system and the other system has a lower learning rate and higher retention rate (i.e. lower forgetting rate) which we refer to as the slow system. For simplicity the same retention and learning rate was used for all objects for each animal.

In the recording session, PV and FV tasks and passage of time were modelled by forgetting rate only as there was no RPE to drive learning (error-clamp period). The behavioral data that was used to fit the model in this period was the FV gaze bias between good and bad objects in the four time points were FV was done (before, immediately after, 20 minutes after and a day after). An exponential function with scaling factor as an additional parameter was used to map values to gaze bias. The choice of exponential function was based on the previous graded-reward studies ^21,51^.

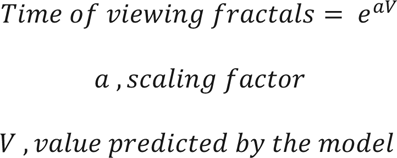

On the other hand, the reversal task was modelled by the full equation including forgetting and learning terms. The behavioral data used for fitting the model in this task was the choice trial performance. In order to map choice rate to value to compare with predicted values of the model a sigmoid function was used in accordance with previous studies with slope of sigmoid as an additional parameter.

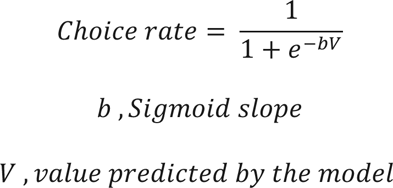

Based on these two models, different combinations of parameters were introduced. In one combination, we assumed different forgetting rates for each task used in the paradigm, that is FC reverse, passive-viewing, free-viewing, irrelevant search tasks and one extra parameter for forgetting in the next day. In another combination, we assumed one forgetting rate for active (FC reverse), one for passive (passive-viewing, free-viewing and irrelevant search) tasks and another for the next day.

Additionally and to account for the possibility that the good and bad objects could become linked in learning such that changes in value of a single good object may provoke changes in value estimate for other unobserved good and/or bad objects, the value update rule was done by taking into account the covariance matrix between object values as a normative approach suggested previously ^52^.

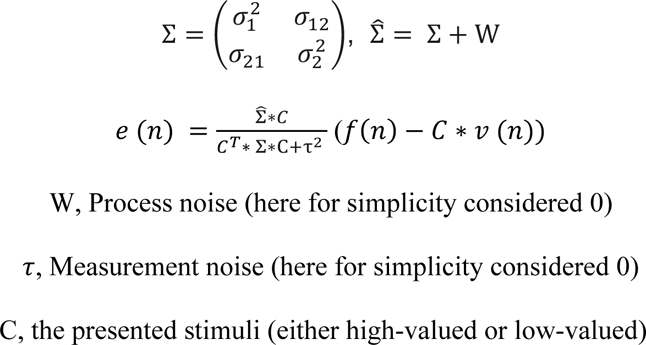

The model prediction evolved throughout the session as the monkey engaged in the experiment, but the ground truth for fitting the model consisted of choice rate in 16 choice trials in FC reversal task and 80 trials in four FV tasks (one before and three after the reversal). FV model fitting was performed using the difference of viewing good vs bad objects.

Sum of squared-error (SSE) was used as cost function to fit the model to behavior. Since there were two different tasks (FC, FV), a multi-objective genetic algorithm (GA) was applied using MATLAB *gamultiobj* function to simultaneously fit both choice rate and time of viewing the fractals in FV tasks as the two cost functions had different scales.

Taken together, eight models were fit to behavioral data. Four single-rate and four double-rate models.

The first single rate model had 6 parameters: two parameters for learning and forgetting in FC reverse, one for slope of the sigmoid function, one for forgetting in trials during passive-viewing (64 trials), free-viewing (20 trials) and irrelevant search (240 trials) tasks during the 20 minutes, one for scaling factor of the exponential function and eventually one parameter for forgetting factor of the next day. Covariance of two (good and bad) distributions (𝑠𝑖𝑔𝑚𝑎) was considered one. In other words, the amount of value update of the seen fractal was applied to the unseen fractal. The second single-rate model had sigma, the covariance of two distributions, as the extra parameter with a total of 7 parameters. The third single-rate model had different parameters for forgetting rates in passive-viewing, free-viewing and irrelevant search with a total of 8 parameters. The fourth single rate model had the same parameters with sigma as an additional parameter and a total of 9 parameters.

The first two-rate model had 10 parameters: four parameters for learning and forgetting rates of slow and fast systems in FC reverse, one for slope of the sigmoid function, two forgetting rates of slow and fast in trials during passive-viewing, free-viewing and irrelevant search during the 20 minutes, one for scaling factor of the exponential function and eventually two parameters for slow and fast forgetting of the next day. The second two-rate model followed the same parameters with sigma as an extra parameter. The third double-rate model had different parameters for forgetting rates of slow and fast systems in passive-viewing, free-viewing and irrelevant search with a total of 14 parameters. Finally, the fourth model had the same parameters with sigma as an additional parameter and a total of 15 parameters.

Model selection was performed based on information criteria methods exploiting both Akaike and Bayesian information criteria (AIC, BIC) obtaining from following formula using SSE:

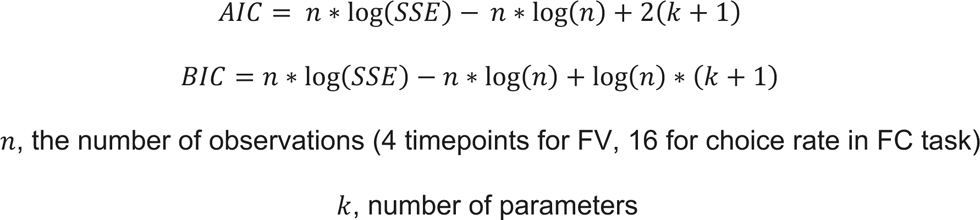

### Statistical analysis

One-sample T-test was used to compare preferred and non-preferred firing rate in neural response, as well as response to good vs bad fractals in LFP power for the averaged early and late epochs, separately. One-way ANOVA test was utilized to examine the effect of memory time points on firing rate and LFP signal. Tukey-HSD test was utilized for subsequent post-hoc analysis. Chi-square test was used to compare the percentage of good-preferring and bad-preferring neurons in each memory time point in AUC distribution. For population-level analysis with pPCA, initially we conducted a resampling procedure. In each iteration, we randomly selected a dataset from the pool of neurons with replacement, ensuring that the dataset’s size matched the total number of neurons. pPCA was then applied to the selected dataset, and this procedure was repeated 100 times. Subsequently, a MANOVA test was performed, using residual PC1 and PC2 data from the 100 datasets to investigate the effect of memory time points on neural states. Eventually, Hotelling’s T^2^ test was used to compare the neural state between two memory periods ^53^

## Supplementary Figures

**Suppl Figure 1.**
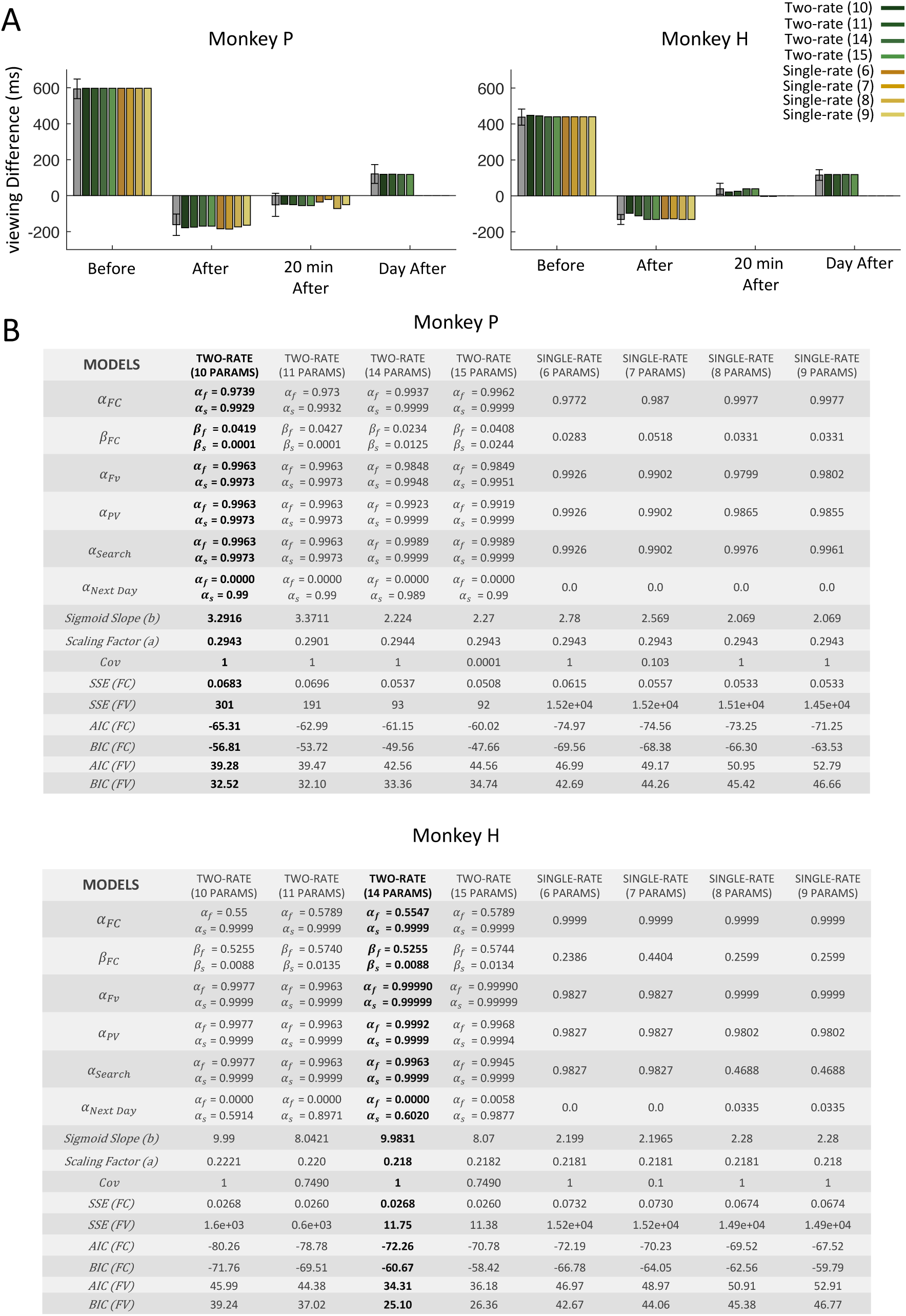
List of various parametrizations for single- and two-rate models and their performance in predicting FV behavior. (A) same format as Fig 1H but for all model parameterizations for each monkey (B) Table of the parameter fits and model selection metrics of each monkey. Sum of squared-errors (SSE) and model selection using Akaike and Bayesian information criteria methods (AIC, BIC) for predicting choice in value training (FC) and gaze bias in (FV) tasks, separately.

**Suppl Figure 2.**
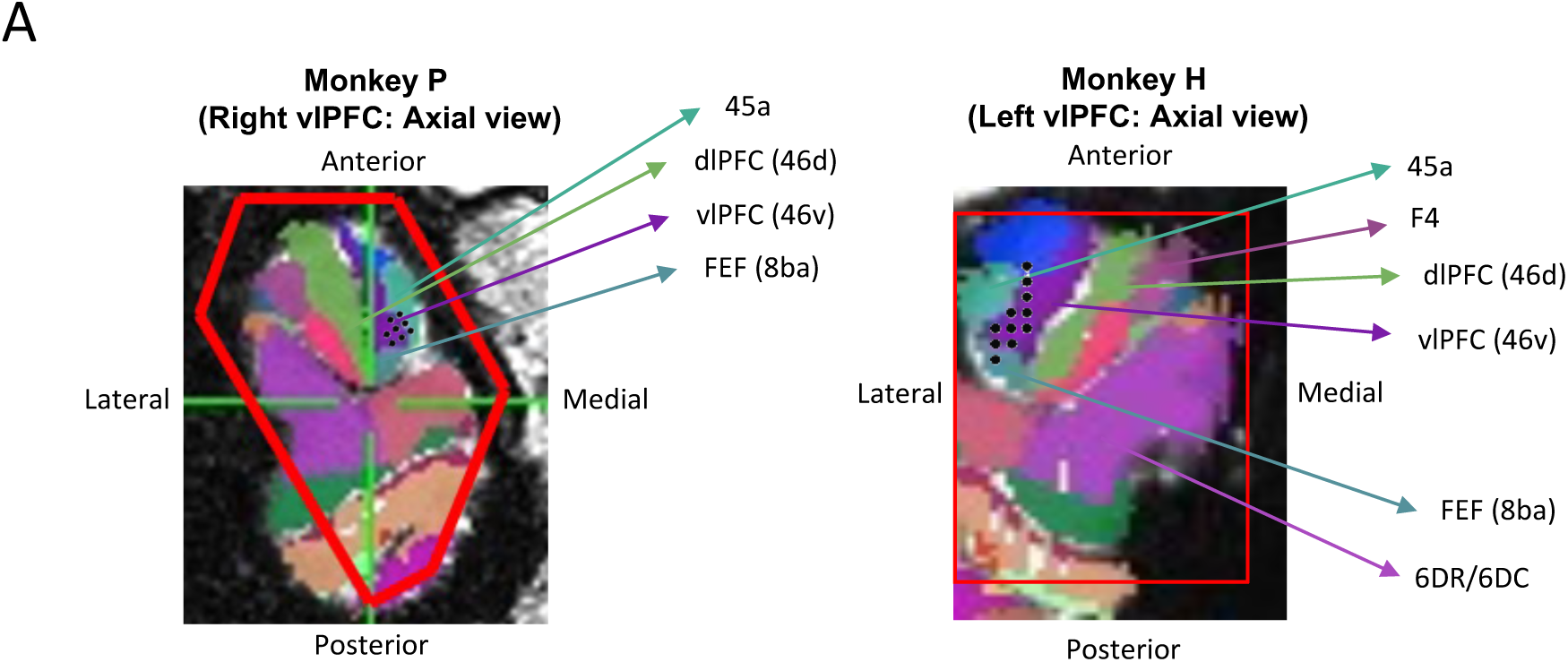
Recording locations in the ventrolateral prefrontal cortex (vlPFC) Axial views showing recorded locations as black dots with NMT atlas warped to native space and region numbers marked and color-coded. The red contour shows the chamber borders in Monkey P (left) and Monkey H (right).

**Suppl Figure 3.**
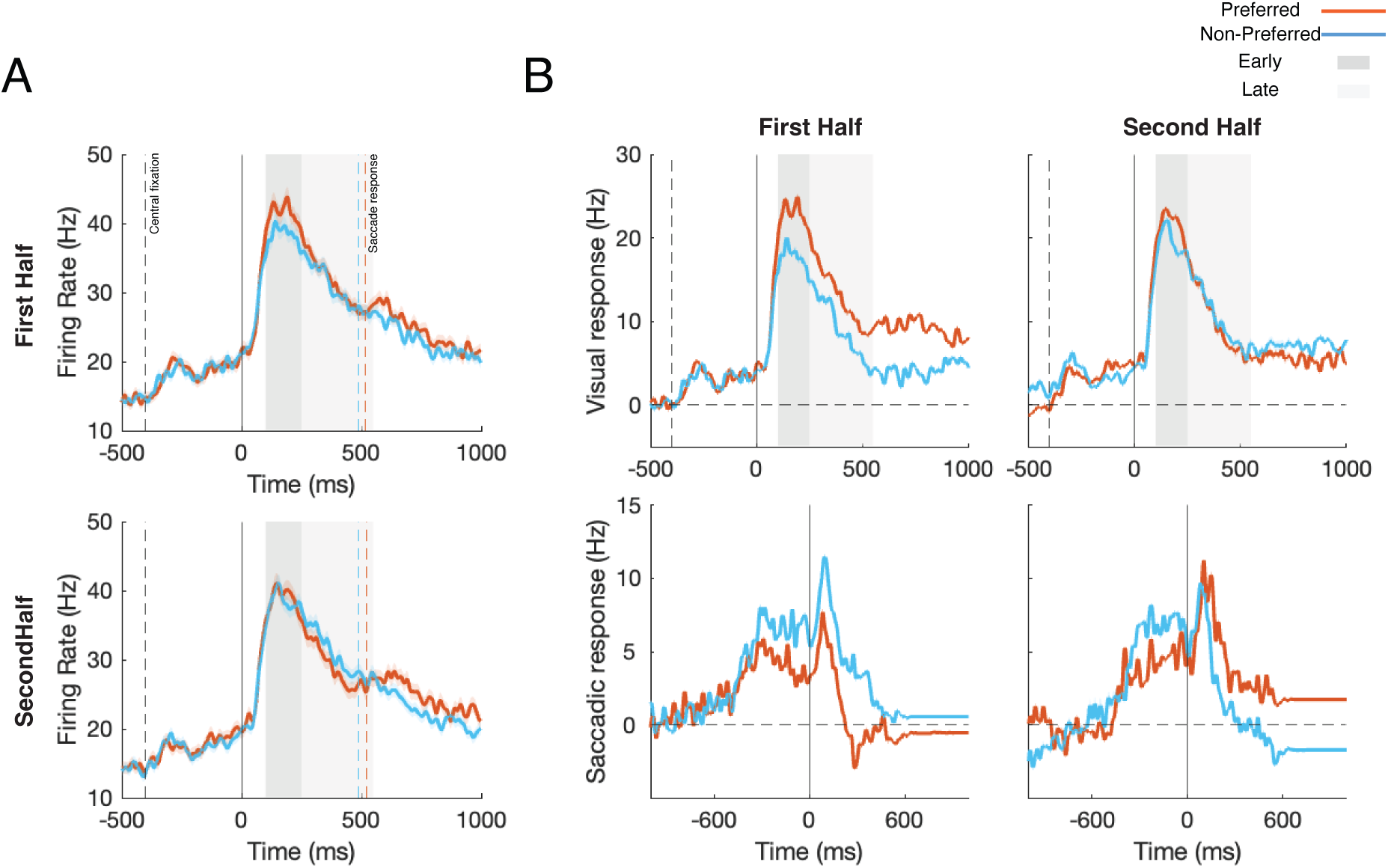
The average neural response during value reversal training. (A) Population average PSTH for preferred and non-preferred values in reversal training task. The reversal learning is divided and averaged across trials in the first and second halves of a block (B) Deconvolution of visual (left) and saccadic (right) parts of average population responses in the first and second halves of the reversal training showing presence of visually evoke and saccade related components.

**Suppl Figure 4.**
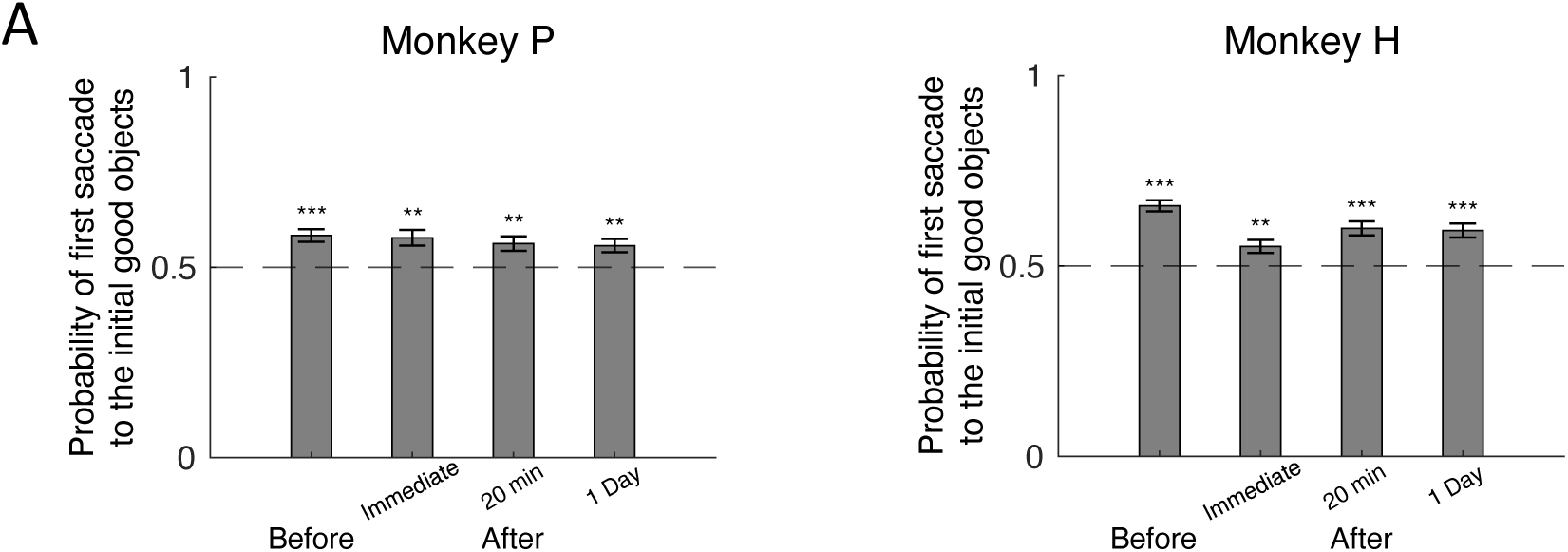
First saccade direction in FV. Probability of first saccade toward the initial good objects across timepoints in FV task for monkey P (left panel) and monkey H (right Panel). The horizontal dashed-line indicates the chance level and the probability above 0.5 means higher probability of making the first saccade toward the initial good objects and vice versa.

**Suppl Figure 5.**
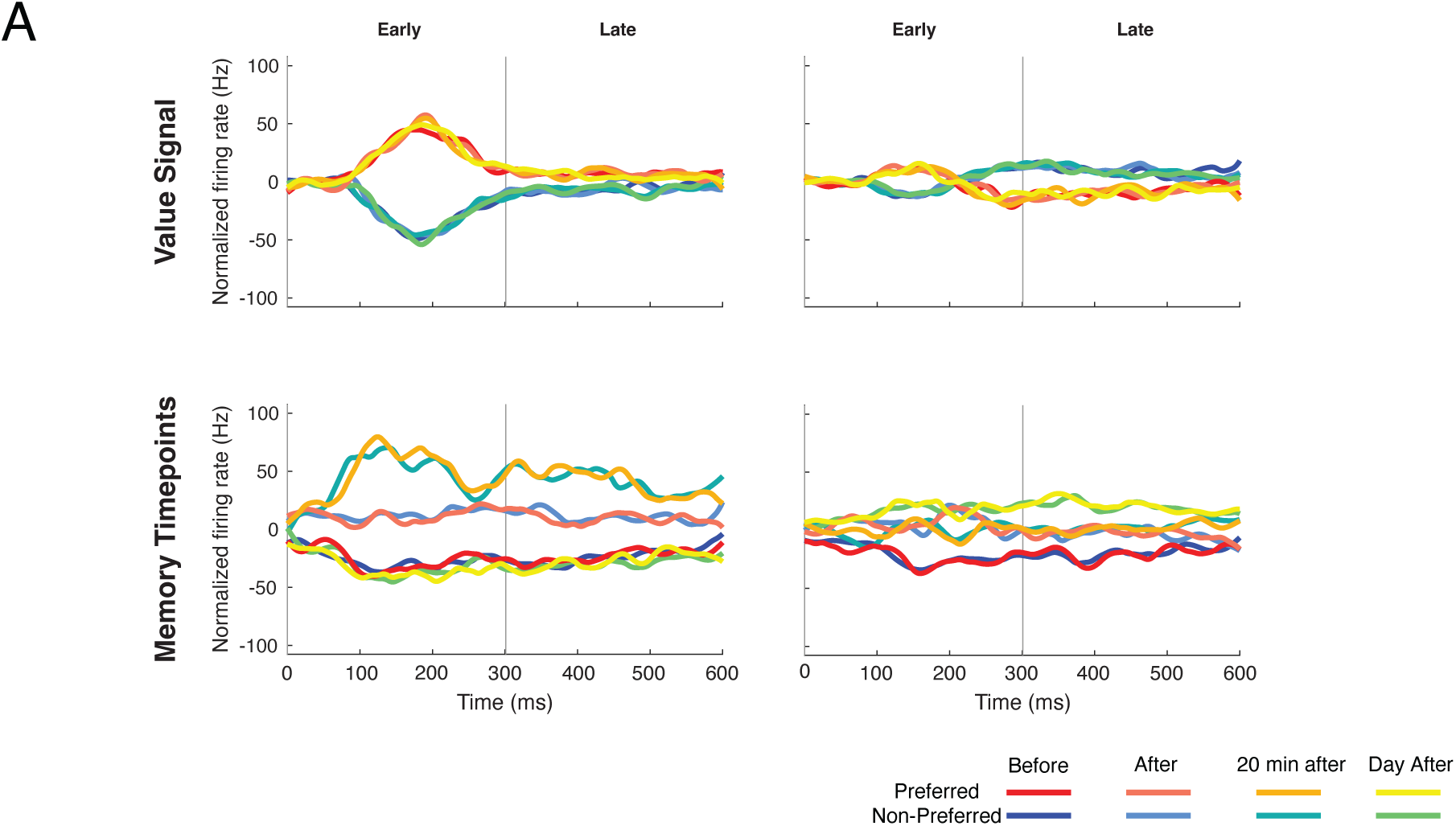
Population state dynamics using demixed-PCA method. (A) Demixed principal components (dPCA). Top row: top two components (explaining 6% and 1.4% of variability, respectively) that are related to value dimension (pref / non-pref); bottom row: top first and third components (explaining 14.3% and 3.5% of variability, respectively) that are related to encoding four time points. In each subplot, the full data are projected onto the respective dPCA decoder axis, so that there are 8 lines corresponding to 8 conditions (two for preference x four time points). Note that consistent with our partial-PCA method, two value components (first row) seem to be showing early and late components. Moreover, the two components in probe time points (bottom row) are similar to PC1 and PC2 in our pPCA results the first one travels and comes back while the second shows a sustained shift.

**Suppl Figure 6.**
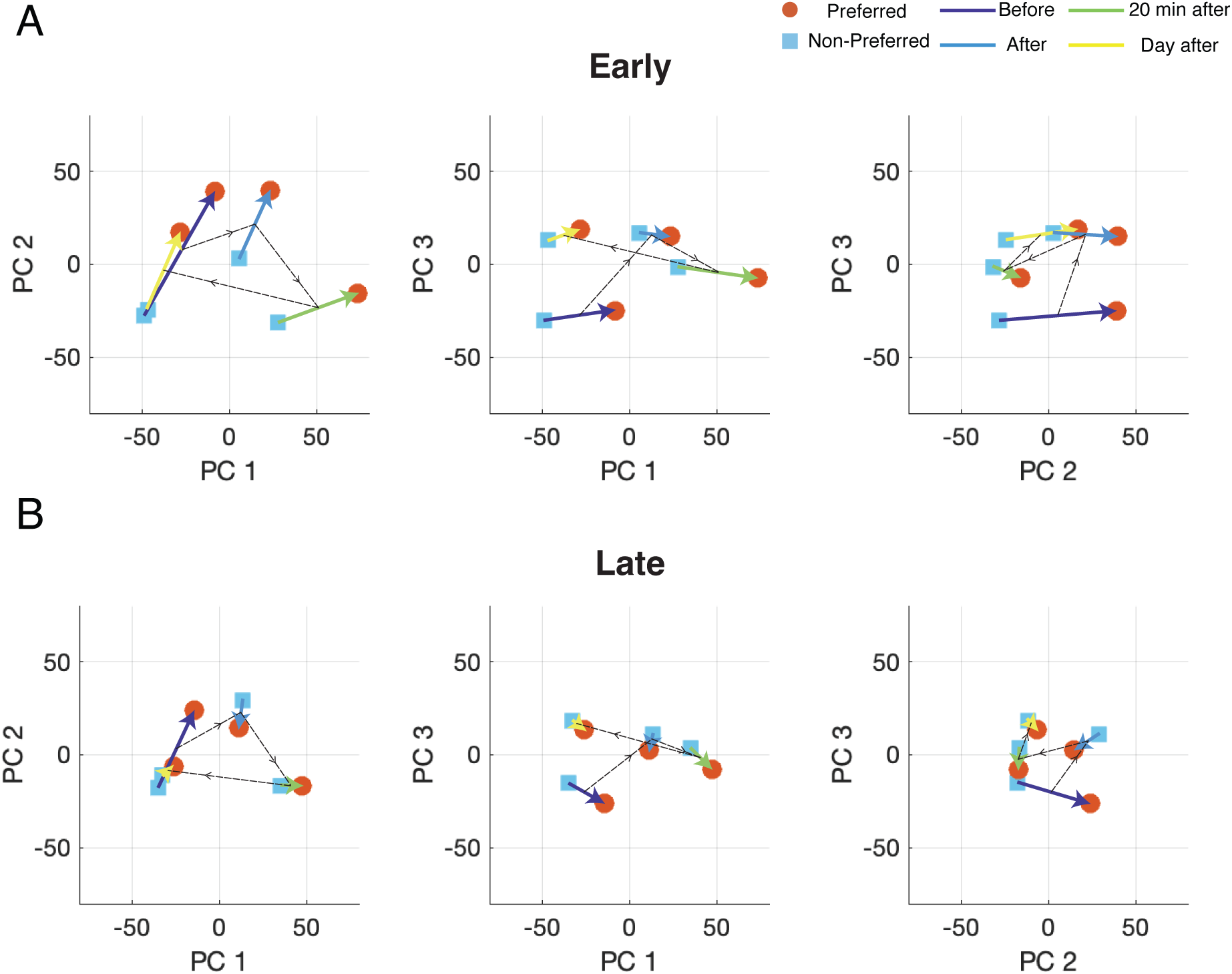
2D projection of population responses across value probes using PCA. same format as in Fig 4 but using normal PCA. Note how PC2 is roughly similar to the value dimension but less optimized compared to p-PCA results.

**Suppl Figure 7.**
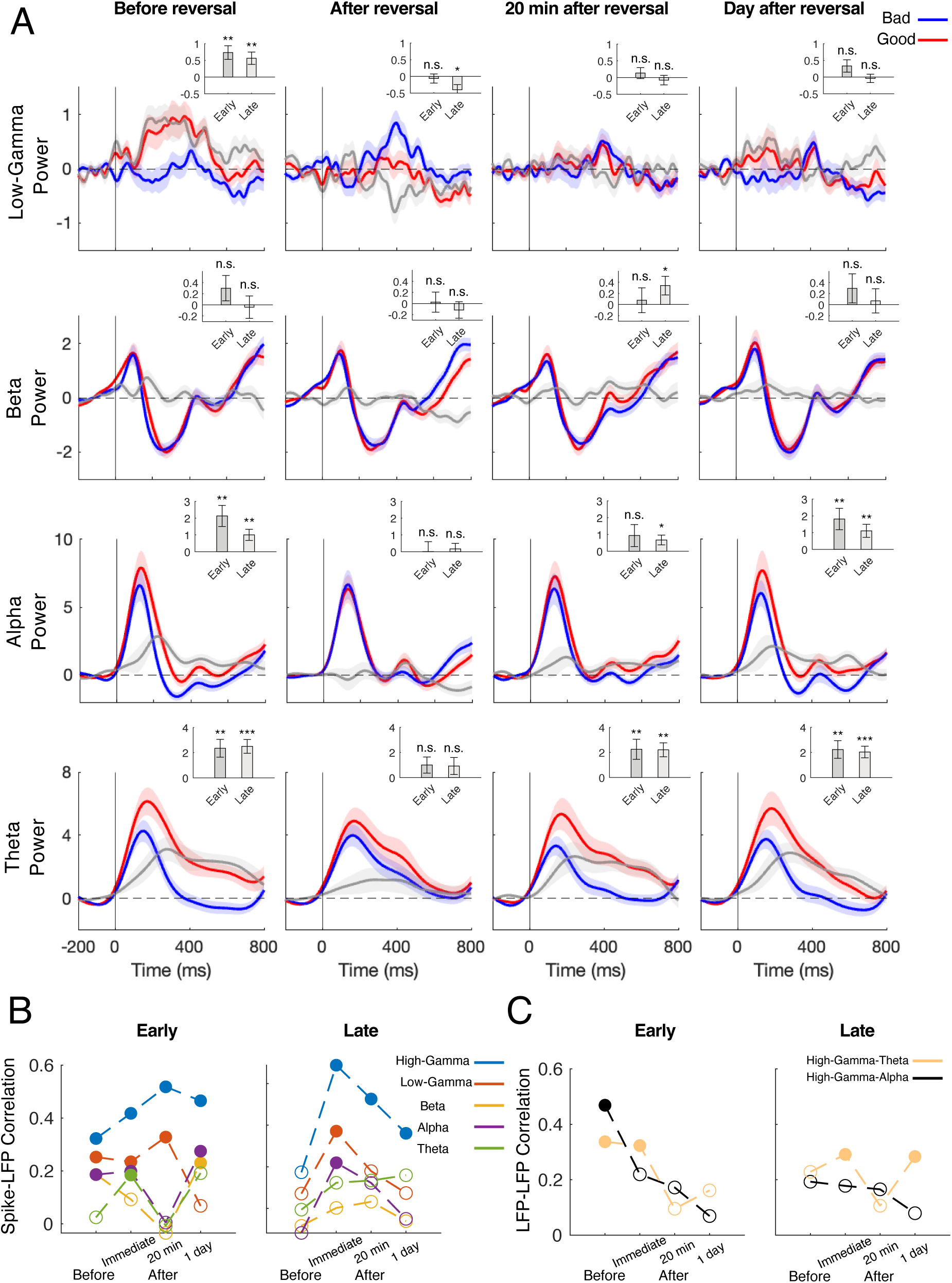
LFP frequency band modulations in time and across value probes. (A) Average low-Gamma (30-60 Hz, top row), Beta (12-30 Hz, second row), Alpha (8-12 Hz, third row) and Theta (4-7 Hz, bottom row) band power in response to good (red) and bad (blue) fractals across memory periods and the response difference (value signal, gray). Inset bar plots indicate average value signal in early and late epochs. (B) correlation between neural value signal and frequency bands across memory periods. Filled circles indicate significant correlation. (C) Correlation between High-Gamma and Alpha and Theta band value power across timepoints. Filled circles indicate a significant correlation.

**Suppl Figure 8.**
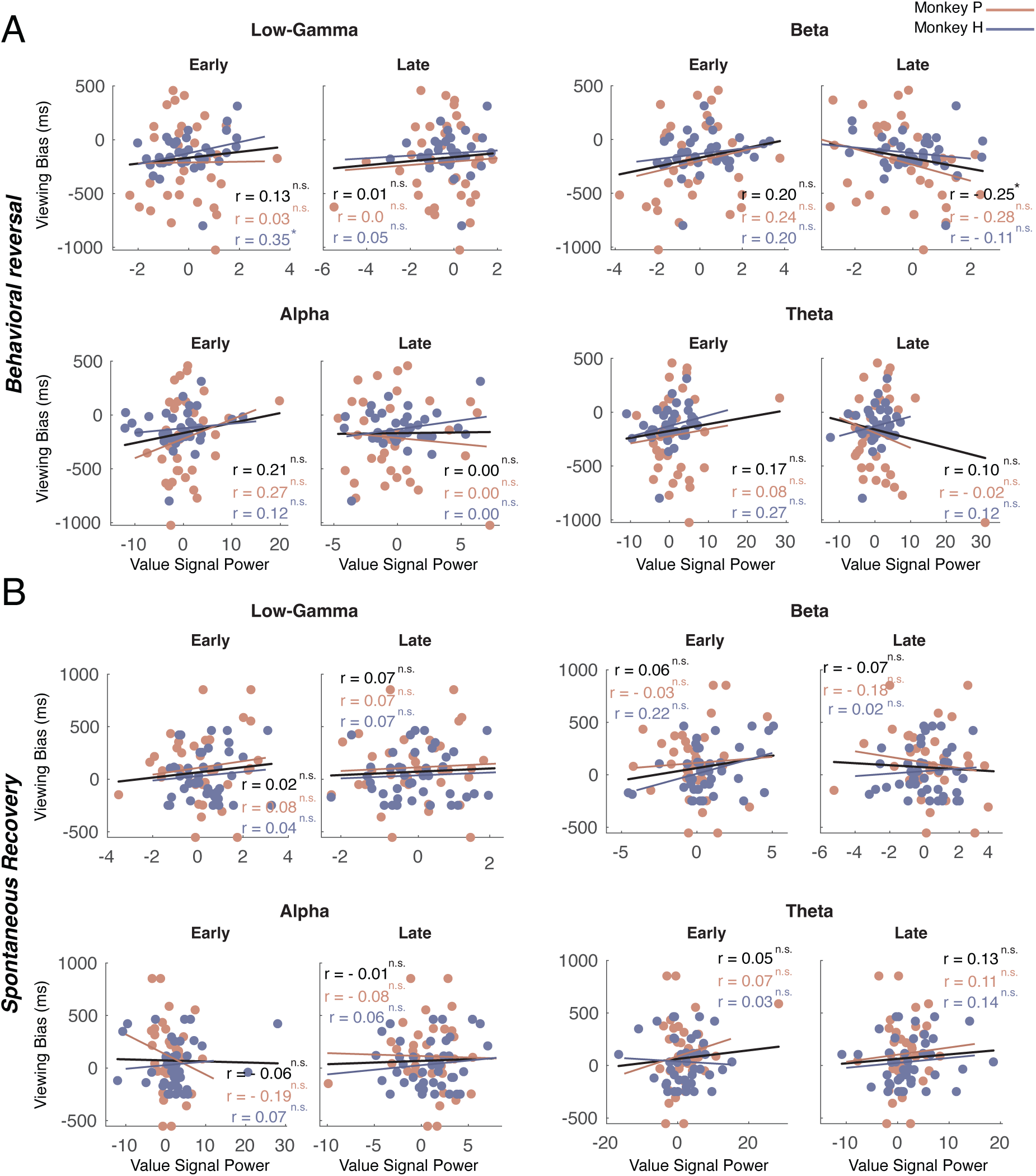
Correlation of LFP bands and gaze bias immediately after and a day after reversal. (A) same format as Fig 5C but for frequency band other than high-Gamma (B) same format as Fig 5D but for frequency band other than high-Gamma

## References

1 Ghazizadeh, A., Griggs, W. & Hikosaka, O. Object-finding skill created by repeated reward experience. Journal of Vision 16, 17–17, doi:10.1167/16.10.17 (2016).

2 Abbaszadeh, M., Panjehpour, A., Amin Alemohammad, S. M., Ghavampour, A. & Ghazizadeh, A. Prefrontal cortex encodes value pop-out in visual search. iScience 26, 107521, doi:10.1016/j.isci.2023.107521 (2023).

3 Abraham, W. C. & Robins, A. Memory retention–the synaptic stability versus plasticity dilemma. Trends in neurosciences 28, 73–78 (2005).

4 Hikosaka, O., Kim, H. F., Yasuda, M. & Yamamoto, S. Basal ganglia circuits for reward value–guided behavior. Annual review of neuroscience 37, 289–306 (2014).

5 Hikosaka, O., Ghazizadeh, A., Griggs, W. & Amita, H. Parallel basal ganglia circuits for decision making. Journal of neural transmission 125, 515–529 (2018).

6 Alexander, G. E., DeLong, M. R. & Strick, P. L. Parallel organization of functionally segregated circuits linking basal ganglia and cortex. Annual review of neuroscience 9, 357–381 (1986).

7 Yin, H. H. & Knowlton, B. J. The role of the basal ganglia in habit formation. Nature Reviews Neuroscience 7, 464–476, doi:10.1038/nrn1919 (2006).

8 Kim, H. F. & Hikosaka, O. Distinct basal ganglia circuits controlling behaviors guided by flexible and stable values. Neuron 79, 1001–1010 (2013).

9 Griggs, W. S. et al. Flexible and stable value coding areas in caudate head and tail receive anatomically distinct cortical and subcortical inputs. Frontiers in neuroanatomy 11, 106 (2017).

10 Bocincova, A., Buschman, T. J., Stokes, M. G. & Manohar, S. G. Neural signature of flexible coding in prefrontal cortex. Proceedings of the National Academy of Sciences 119, e2200400119 (2022).

11 Miller, E. K. The prefontral cortex and cognitive control. Nature Reviews Neuroscience 1, 59–65, doi:10.1038/35036228 (2000).

12 Warden, M. R. & Miller, E. K. Task-Dependent Changes in Short-Term Memory in the Prefrontal Cortex. The Journal of Neuroscience 30, 15801–15810, doi:10.1523/jneurosci.1569-10.2010 (2010).

13 Barraclough, D. J., Conroy, M. L. & Lee, D. Prefrontal cortex and decision making in a mixed-strategy game. Nat Neurosci 7, 404–410, doi:10.1038/nn1209 (2004).

14 Ghazizadeh, A., Hong, S. & Hikosaka, O. Prefrontal cortex represents long-term memory of object values for months. Current Biology 28, 2206–2217. e2205 (2018).

15 Borra, E., Gerbella, M., Rozzi, S. & Luppino, G. Projections from caudal ventrolateral prefrontal areas to brainstem preoculomotor structures and to basal ganglia and cerebellar oculomotor loops in the macaque. Cerebral Cortex 25, 748–764 (2015).

16 Middleton, F. A. & Strick, P. L. Basal-ganglia ‘projections’ to the prefrontal cortex of the primate. Cerebral cortex 12, 926–935 (2002).

17 Kim, H. F., Ghazizadeh, A. & Hikosaka, O. Dopamine neurons encoding long-term memory of object value for habitual behavior. Cell 163, 1165–1175 (2015).

18 Yasuda, M., Yamamoto, S. & Hikosaka, O. Robust Representation of Stable Object Values in the Oculomotor Basal Ganglia. The Journal of Neuroscience 32, 16917–16932, doi:10.1523/jneurosci.3438-12.2012 (2012).

19 Smith, M. A., Ghazizadeh, A. & Shadmehr, R. Interacting adaptive processes with different timescales underlie short-term motor learning. PLoS biology 4, e179 (2006).

20 Ghazizadeh, A., Griggs, W., Leopold, D. A. & Hikosaka, O. Temporal–prefrontal cortical network for discrimination of valuable objects in long-term memory. Proceedings of the National Academy of Sciences 115, E2135–E2144 (2018).

21 Ghazizadeh, A., Griggs, W. & Hikosaka, O. Ecological origins of object salience: Reward, uncertainty, aversiveness, and novelty. Frontiers in neuroscience 10, 378 (2016).

22 Kobak, D. et al. Demixed principal component analysis of neural population data. elife 5, e10989 (2016).

23 Buzsaki, G. & Draguhn, A. Neuronal oscillations in cortical networks. science 304, 1926–1929 (2004).

24 Palva, S. & Palva, J. M. Functional roles of alpha-band phase synchronization in local and large-scale cortical networks. Frontiers in psychology 2, 204 (2011).

25 Belitski, A. et al. Low-frequency local field potentials and spikes in primary visual cortex convey independent visual information. Journal of Neuroscience 28, 5696–5709 (2008).

26 Khawaja, F. A., Tsui, J. M. & Pack, C. C. Pattern motion selectivity of spiking outputs and local field potentials in macaque visual cortex. Journal of Neuroscience 29, 13702–13709 (2009).

27 Einevoll, G. T., Kayser, C., Logothetis, N. K. & Panzeri, S. Modelling and analysis of local field potentials for studying the function of cortical circuits. Nature Reviews Neuroscience 14, 770–785, doi:10.1038/nrn3599 (2013).

28 Kahneman, D. Thinking, fast and slow. (macmillan, 2011).

29 Brandon, T. H., Vidrine, J. I. & Litvin, E. B. Relapse and relapse prevention. Annu. Rev. Clin. Psychol. 3, 257–284 (2007).

30 Hunt, W. A., Barnett, L. W. & Branch, L. G. Relapse rates in addiction programs. Journal of clinical psychology (1971).

31 Medina, J. F., Garcia, K. S. & Mauk, M. D. A mechanism for savings in the cerebellum. Journal of Neuroscience 21, 4081–4089 (2001).

32 Coltman, S. K., Cashaback, J. G. A. & Gribble, P. L. Both fast and slow learning processes contribute to savings following sensorimotor adaptation. J Neurophysiol 121, 1575–1583, doi:10.1152/jn.00794.2018 (2019).

33 Brashers-Krug, T., Shadmehr, R. & Bizzi, E. Consolidation in human motor memory. Nature 382, 252–255 (1996).

34 Thoroughman, K. A. & Shadmehr, R. Electromyographic correlates of learning an internal model of reaching movements. Journal of Neuroscience 19, 8573–8588 (1999).

35 Sun, X. et al. Cortical preparatory activity indexes learned motor memories. Nature 602, 274–279, doi:10.1038/s41586-021-04329-x (2022).

36 Van Kerkoerle, T. et al. Alpha and gamma oscillations characterize feedback and feedforward processing in monkey visual cortex. Proceedings of the National Academy of Sciences 111, 14332–14341 (2014).

37 Dom, G., Sabbe, B., Hulstijn, W. & van Den Brink, W. Substance use disorders and the orbitofrontal cortex: Systematic review of behavioural decision-making and neuroimaging studies. The British Journal of Psychiatry 187, 209–220, doi:10.1192/bjp.187.3.209 (2005).

38 Schoenbaum, G. & Shaham, Y. The role of orbitofrontal cortex in drug addiction: a review of preclinical studies. Biol Psychiatry 63, 256–262, doi:10.1016/j.biopsych.2007.06.003 (2008).

39 Davidson, B. et al. Deep brain stimulation of the nucleus accumbens in the treatment of severe alcohol use disorder: a phase I pilot trial. Molecular Psychiatry 27, 3992–4000, doi:10.1038/s41380-022-01677-6 (2022).

40 Mani, P., Kelley, V. & Alexander, Y. Nucleus Accumbens and Its Role in Reward and Emotional Circuitry: A Potential Hot Mess in Substance Use and Emotional Disorders. AIMS Neuroscience 4, 52–70, doi:10.3934/Neuroscience.2017.1.52 (2017).

41 Kennerley, S. W., Walton, M. E., Behrens, T. E., Buckley, M. J. & Rushworth, M. F. Optimal decision making and the anterior cingulate cortex. Nature neuroscience 9, 940–947 (2006).

42 Wallis, J. D. Orbitofrontal cortex and its contribution to decision-making. Annu. Rev. Neurosci. 30, 31–56 (2007).

43 Paton, J. J., Belova, M. A., Morrison, S. E. & Salzman, C. D. The primate amygdala represents the positive and negative value of visual stimuli during learning. Nature 439, 865–870 (2006).

44 Kim, C., Kroger, J. & Kim, J. A functional dissociation of conflict processing within anterior cingulate cortex. Nature Precedings, doi:10.1038/npre.2008.2505.1 (2008).

45 Isoda, M. & Hikosaka, O. Switching from automatic to controlled action by monkey medial frontal cortex. Nature neuroscience 10, 240–248 (2007).

46 Tabu, H., Mima, T., Aso, T., Takahashi, R. & Fukuyama, H. Functional relevance of pre-supplementary motor areas for the choice to stop during Stop signal task. Neuroscience research 70, 277–284 (2011).

47 Aron, A. R. & Poldrack, R. A. Cortical and subcortical contributions to stop signal response inhibition: role of the subthalamic nucleus. Journal of Neuroscience 26, 2424–2433 (2006).

48 Nadian, M. H., Farmani, S. & Ghazizadeh, A. A novel methodology for exact targeting of human and non-human primate brain structures and skull implants using atlas-based 3D reconstruction. Journal of Neuroscience Methods 391, 109851 (2023).

49 Ghazizadeh, A., Fields, H. L. & Ambroggi, F. Isolating event-related neuronal responses by deconvolution. Journal of neurophysiology 104, 1790–1802 (2010).

50 Cohen, M. X. Analyzing neural time series data: theory and practice. (MIT press, 2014).

51 Ghazizadeh, A. & Hikosaka, O. Common coding of expected value and value uncertainty memories in the prefrontal cortex and basal ganglia output. Science advances 7, eabe0693 (2021).

52 Dayan, P. & Jyu, A. Uncertainty and Learning. IETE Journal of Research 49, 171–181, doi:10.1080/03772063.2003.11416335 (2003).

53 Trujillo-Ortiz, A. HotellingT2, <https://www.mathworks.com/matlabcentral/fileexchange/2844-hotellingt2> (2005).

